# Evolution of resistance breaking in spinach downy mildew revealed through pangenome graph analysis

**DOI:** 10.1101/2025.05.23.655769

**Authors:** Petros Skiadas, Melanie N. Mendel, Joyce Elberse, Guido Van den Ackerveken, Ronnie de Jonge, Michael F. Seidl

## Abstract

**Background:** Filamentous plant pathogens secrete effectors to successfully establish host infections. In resistant crop varieties, plant immunity can be triggered by immune receptors that recognize these effectors. Resistant crop varieties are grown in large-scale monocultures imposing strong selection pressure on pathogens, driving rapid evolution of effector repertoires resulting in the frequent breakdowns of resistance within just a few growing seasons. The oomycete *Peronospora effusa*, responsible for downy mildew on spinach, is an example of a rapidly adapting pathogen, but it is yet unknown how *P. effusa* can successfully overcome resistance of spinach by genomic adaptations.

**Results:** To close this knowledge gap, we here generated genome assemblies and constructed a pangenome graph for 19 isolates corresponding to 19 officially denominated resistance-breaking *P. effusa* races, which can cause disease on a differential set of spinach cultivars. Haplotype-resolved pangenome graph analyses revealed that many isolates emerged from recent sexual recombination, yet others evolved via prolonged asexual reproduction and loss of heterozygosity. By phasing effector candidates to determine their allelic variation, we identified effector candidates associated to resistance breaking of spinach varieties and reconstructed the evolutionary events that led to their diversification.

**Conclusions:** The here developed and applied computational genomics approaches offer unparalleled insights into the molecular mechanisms of the rapid evolution of *P. effusa*, and points to potential targets for future resistance breading.

## Background

To establish a successful infection, filamentous plant pathogens secrete so called effector molecules that promote host colonisation by, for example, circumventing or suppressing host immune responses [1,2]. Plant immunity can be activated in a gene-for-gene manner, where a single plant resistance protein is activated in response to a single pathogen effector, known by the term “avirulence factor”, leading to a strong hypersensitive response that stops pathogen infection, a process termed effector-triggered immunity [3,4]. In turn, pathogens that do not express, have mutations in, or have lost this particular effector can break the resistance of the host plant and thereby reestablish their virulence [5,6].

Effector-triggered immunity is qualitative, often monogenic or oligogenic, and thus it is a desirable trait for the development of resistant crop cultivars. In agriculture, these resistant cultivars are often deployed in large monocultures [7]. However, this agricultural practice exerts strong selective pressure on pathogens to overcome resistances, leading to the rapid diversification of pathogen populations [8,9]. On the genomic level, this strong selection drives the emergence of novel or the diversification of existing effectors to avoid host recognition and to re-establish successful host colonization [1]. Unsurprisingly, host resistances are often broken within only a few growing seasons after the introduction of resistant cultivars in the fields [10].

Oomycetes are a diverse and widespread group of filamentous organisms including many important pathogens of plants, causing devastating diseases and resulting in severe damage in agriculture and natural ecosystems [11,12]. To establish a successful infection, oomycetes utilize two classes of effector proteins: apoplastic effectors, which act outside the plant cells, and cytoplasmic effectors, which are translocated inside the plant cell [13]. Two main families of cytoplasmic effectors have been characterized in oomycetes thus far, the RXLR and Crinkler (or CRN) effectors [14,15]. These two effector families are characterized by the presence of conserved motifs at the N-terminus downstream of the signal peptide, which have been hypothesized to contribute to effector translocation into the host cell or to their secretion [16–19]. The C-terminal regions of these effectors vary significantly and are responsible for the effectors’ functions in the plant cell [13,18,20]. This region can be recognized by hosts’ immune systems, triggering hypersensitive response and host resistances [21]. Several nucleotide-binding leucine-rich repeat receptors (NLRs) have been identified that can recognise avirulence proteins, including the R3a from a wild potato species that recognises the AVR3a effector of the oomycete tomato and potato late blight pathogen *Phytophthora infestans* [21,22]. However, aside from a few effectors characterized in *Phytophthora*, *Hyaloperonospora*, and *Plasmopara* species, little is known about effector diversity in other oomycetes.

The obligate biotrophic oomycete *Peronospora effusa* causes downy mildew on spinach, the economically most important disease of cultivated spinach worldwide [23,24]. This pathogen has been traditionally managed by fungicides and the extensive deployment of genetic disease resistances [25,26]. Resistant spinach cultivars, for example those with the NLR *RPF1*, are thought to encode receptors that can recognize specific *P. effusa* effectors and thereby induce effector triggered immunity [27]. These cultivars have been extensively used for spinach production and have been the most effective management tool for downy mildew, especially in organic practices [25]. However, *P. effusa* rapidly breaks resistances of newly introduced spinach varieties [24]. Due to this rapid evolution, a new *P. effusa* race is denominated every year based on isolates having the capacity to break spinach resistances, with 20 races denominated thus far [23,27,28]. Like many oomycetes, *P. effusa* is diploid and can reproduce both asexually and sexually, and different isolates show various levels of heterozygosity [29]. Sexual recombination was suggested to be a powerful driver of the emergence of new *P. effusa* races [29,30]. However, we currently lack direct genomic evidence of recombination between the isolates or for the role of heterozygosity in the variation of the effector repertoires between *P. effusa* isolates.

*P. effusa* is one of the few oomycete species with multiple isolates having chromosome-level reference genome assemblies publicly available [31–33]. Additionally, we have recently developed a pangenome approach for in depth comparisons of multiple genome assemblies and their structural annotations in depth [31]. The genome assembly of *P. effusa* is 57.8-60.5 Mb in size, is highly repetitive (56% repeat content), and is organised in 17 core chromosomes, with few isolates having a single additional accessory chromosome with unknown function [31]. On average, of the 10,312 annotated genes, 472 are effector candidates that are often under positive selection and are highly variable between isolates [31]. While it is conceivable that effector variation between the *P. effusa* races is linked to the breakage of spinach resistances, no effector candidates associated with (a)virulence have been identified thus far. Here, we capitalized on our previously developed pangenome framework [31] to analyse chromosome-level genome assemblies for 19 denominated resistance-breaking *P. effusa* isolates. Many *P. effusa* isolates are the result of recent recombination yet few also display patterns of prolonged asexual reproduction and loss of heterozygosity. Importantly, phasing of all effector candidates enabled us to identify specific effectors that can be associated to the breaking of spinach resistance and enabled us to reconstruct the molecular origins of their genomic variation. This provides invaluable insights into the mechanisms behind the rapid evolution of *P. effusa* and highlights targets for future experimental analysis for resistance breading.

## Results

### *Peronospora effusa* genomic variation is due to the expansion of repeats and gene copy numbers

To explore the genomic variation in *P. effusa*, we selected 19 isolates representing 19 races denominated for their capacity to break different combination of resistance alleles introduced in spinach (Figure 1A) [34]. Like other oomycetes, *P. effusa* has both a sexual and an asexual reproduction cycle [29]. To better represent this complex phylogeny, we determined the relationship between the 19 isolates with a neighbor-net phylogenetic network analysis using 200,934 biallelic single nucleotide polymorphisms (SNPs). Next to few clusters of closely related isolates, we observed that isolates that are branching from the centre of the phylogenetic network are likely the result of a recent recombination between distant isolates (e.g. *Pe7*, *Pe11*, or *Pe13;* PHI-test <0.0001 statistically supports recombination between isolates). Moreover, most isolates have long branches due to a high number of unique SNPs (on average 6,605 SNPs or 3.9% of the SNPs are unique per isolate), suggesting prolonged asexual reproduction or recombination with closely related *P. effusa* genotypes that have not been isolated and analysed (Figure 1B).

**Figure 1.**
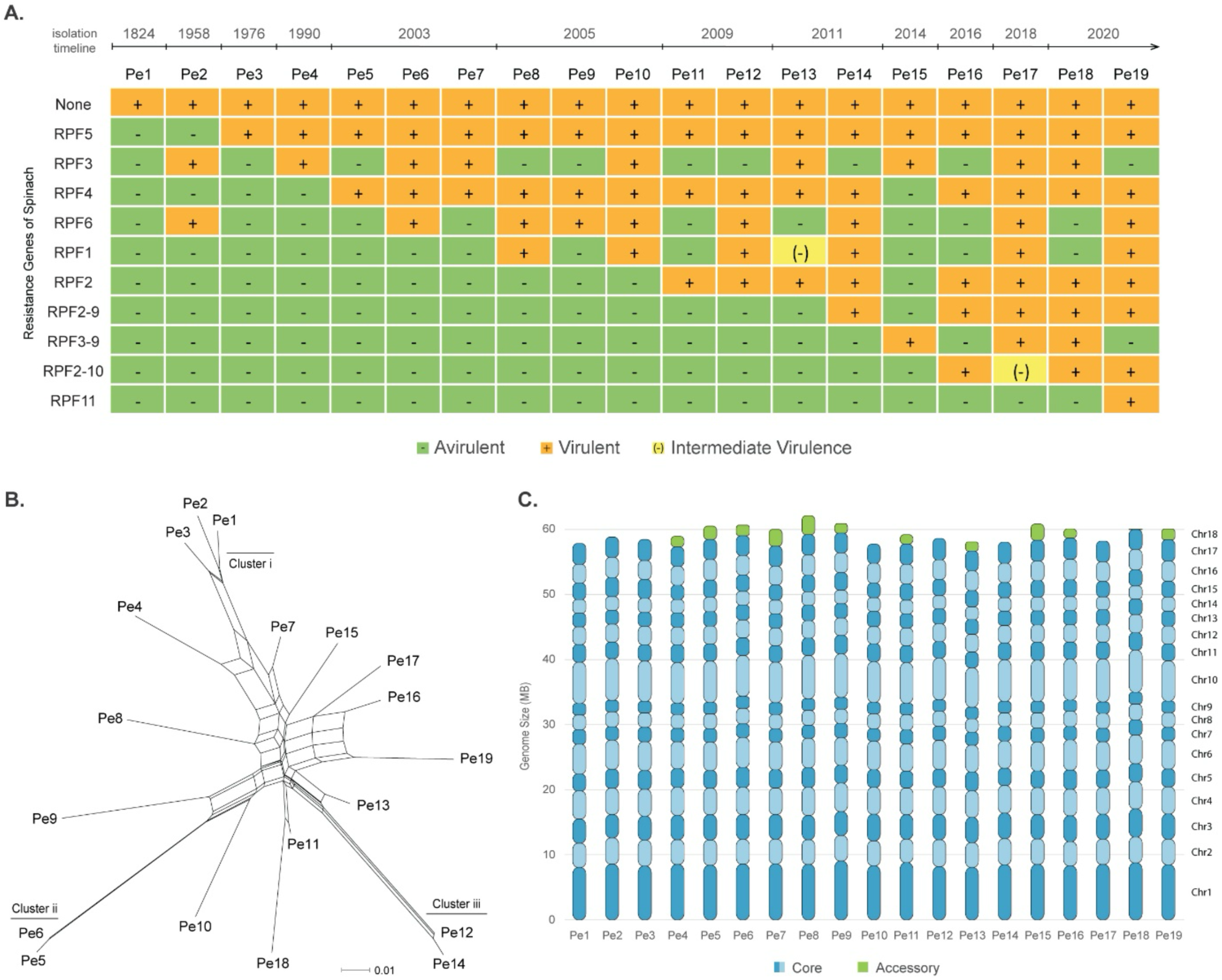
*Peronospora effusa* isolates break various spinach resistances. **A.** The (a)virulence phenotypes of 19 *P. effusa* isolates that break different combination of spinach resistance genes. The year of identification is indicated as well as the NLR resistance genes present in the spinach cultivars that are successfully infected by the virulent *P. effusa* race. **B.** Neighbor-net phylogenetic network indicates the relationships between *P. effusa* isolates. Branch lengths are proportional to the calculated number of substitutions per site. The parallel edges connecting different isolates indicate conflicting phylogenetic signals, suggesting recombination between isolates. **C.** Stacked bar plots display the different chromosomes and their respective sizes for each *P. effusa* isolate. The presence and size of the accessory chromosome 18 is indicated in green.

Of the 19 *P. effusa* isolates, genomes of six have been previously assembled, which revealed 17 highly conserved and collinear core chromosomes and an additional 18^th^ accessory chromosome that is present in few isolates [31]. Here, we generated genome assemblies for the remaining 13 isolates using Nanopore and Illumina sequencing data and created chromosome-level, haploid genome assemblies with total genome assembly sizes ranging between 57.8 and 62.1 Mb (Figure 1C) (Table S1, S2). Out of the 19 genome assemblies, twelve have contigs that share similarity with the previously described accessory chromosome [31], while seven have only contigs that are similar to the 17 core chromosomes. All contigs that match the 17 core chromosomes have telomeric repeats with the repeat motif ‘TTTAGGG’ on both ends, suggesting that we successfully obtained complete and chromosome-level genome assemblies for all 19 isolates. The assembly completeness, evaluated by Benchmarking Universal Single-Copy Ortholog (BUSCO) analysis using the Stramenopiles database (v. odb10), revealed a 99% BUSCO completeness score, which is comparable with previous chromosome-level *P. effusa* genome assemblies [31,35], indicating that these assemblies are highly contiguous and successfully captured most protein-coding regions.

To evaluate the collinearity of the 17 core chromosomes, we performed whole-genome alignments of the 19 genomes based on sequence similarity and on relative position of protein-coding genes. The genome assemblies are nearly completely co-linear, except for few intrachromosomal rearrangements on chromosomes 1, 3, 10, and 17 (Figure S1). These findings corroborate previous observations that the core chromosomes of *P. effusa* are conserved with no gross chromosomal rearrangements [31,33].

To systematically describe the genomic variation of these 19 *P. effusa* isolates, we utilised our previously developed pangenomic approach to directly compare the chromosome-level assemblies without possible reference biases [31]. We used all 19 assemblies to create 17 sequence-resolved pangenome graphs, one for each core chromosome, based on Minigraph-Cactus [36]. The combined pangenome graph has 3,570,190 nodes and 4,858,391 edges, which is only a small fraction of all theoretically possible connections between the nodes, indicating that the addition of genomes does not lead to a complex pangenome structure. Most nodes have only two connections, indicating a mostly linear graph, while only 82 nodes have a degree of ten or higher (Figure S2A). The total size of the combined graph is 104.8 Mb, 80% larger than the average *P. effusa* genome assembly (58 Mb), suggesting that while most of the genome organization is conserved, the pangenome graph nevertheless captures a significant amount of accessory genomic regions (Figure 2A).

**Figure 2.**
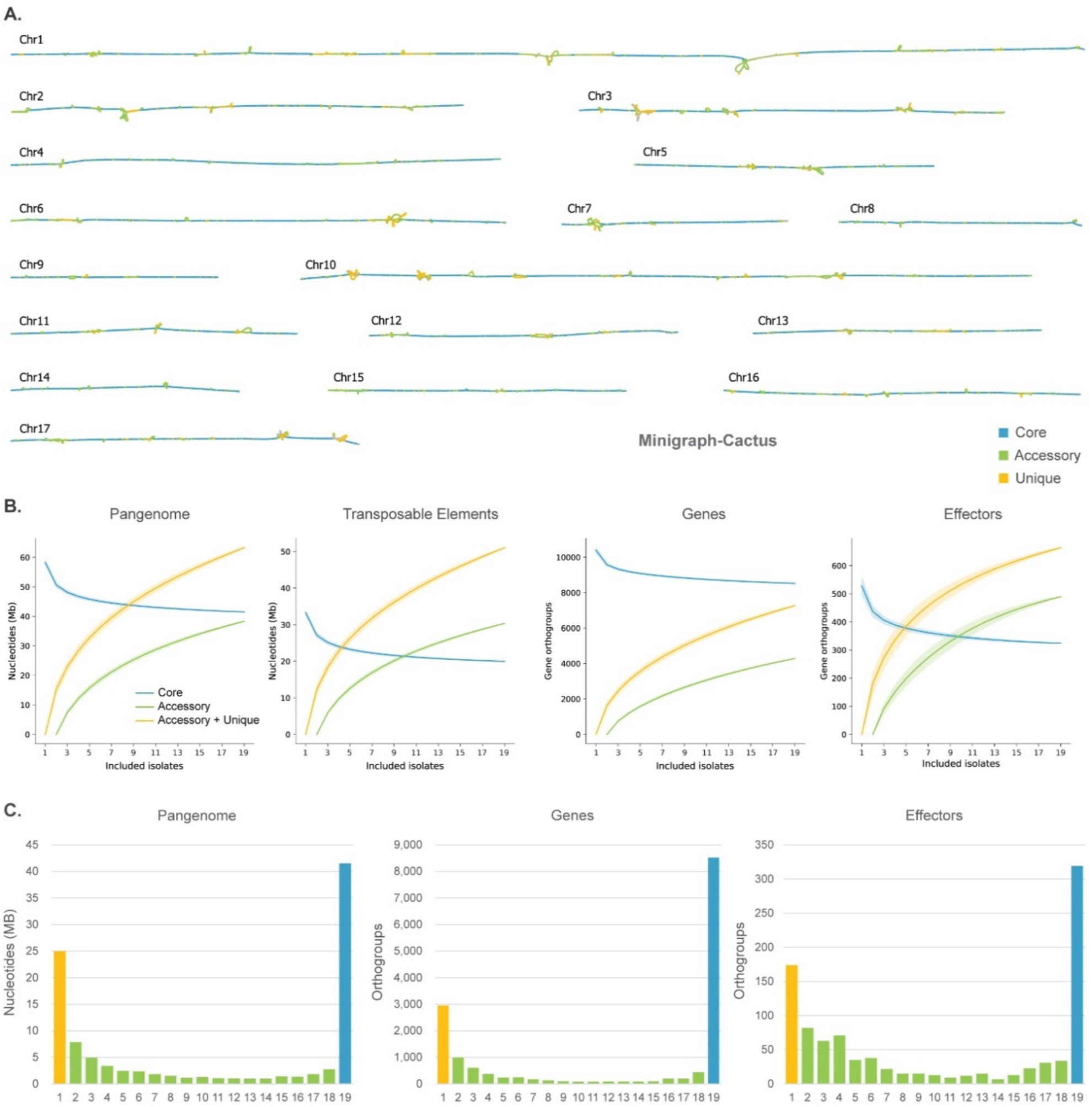
Pangenome graph analysis for 19 *Peronospora effusa* isolates. **A.** Pangenome graphs of the 17 core chromosomes that are present in all *P. effusa* isolates were created with Minigraph-Cactus and visualised with Bandage [36,37]. **Β.** Saturation plots based on the pangenome graph on the nucleotide level for the whole genome and transposable elements, as well as on the gene orthogroup level for all genes and effector candidates. **C.** Bar plot of the total size of pangenome graph nodes that belong to one to 19 isolates separated for all nodes, for nodes that overlap with all gene annotations, and for nodes that overlap with effector gene annotations.

Consistent structural annotation is essential to discover and describe genomic variation between isolates. Following our previous methods, we used a transposable element (TE) library, based on multiple *P. effusa* isolates, to annotate TEs, resulting in 54-57% of each genome being annotated as TEs (Table S3) [31]. For protein-coding gene annotation, we took advantage of our previously developed joined structural annotation approach [31], which resulted in the annotation of 9,571 to 10,540 protein-coding genes for each isolate. The joined structural annotation added between 239 to 866 genes (2.4-9.0%) that have been missed by the conventional approach that annotated each isolate independently. We assigned these genes to orthologous groups based on their relative position on the pangenome graph, creating 15,740, with each isolate being either absent or represented by a single gene in each group. To identify effector candidates, we then searched the predicted protein-coding genes and additional open reading frames for those encoding proteins with a predicted signal peptide and for RXLR and CRN amino acid motifs, which yielded 354 to 475 putative RXLR and 33 to 66 putative CRN effectors per isolate (Table S3), abundances similar to those in previously assembled *P. effusa* isolates [31,33].

The overall genomic variation between the nineteen *P. effusa* isolates can be uncovered by directly querying the pangenome graph. This analysis revealed that 39.6% (around 41.5 Mb) of the *P. effusa* pangenome is conserved, 36.6% (38.3 Mb) is found in two or more isolates, and 23.8% (24.9 Mb) is unique for single isolates (Figure 2A, S2C). Thus, for each isolate on average 71.2% of the genome is core, 26.5% is accessory, and 2.2% is unique (Figure S2C). When we compared this to the previously generated pangenome based on only six isolates [31], we observed that the percentage of core regions had a small decrease (by 9.5%), the accessory regions greatly increased (49.4%), while the unique regions decreased (37.2%). This was expected, since the core regions of six isolates remain largely conserved in the 19 isolates, while the additional isolates add context to the shared variation between *P. effus*a isolates, overall resulting in the unique regions to decrease in abundance and the accessory to increase. Unexpectedly though, the unique regions per isolate still represent a significant portion of their genomes (1.6-4.1%), causing the abundance of unique regions in the 19 isolate pangenome to double in comparison to the six-isolate pangenome, suggesting that there is significant genomic variation yet to be discovered (Figure S2C).

The abundance of genomic variation is also visible in the saturation plots, which reveal an open pangenome in *P. effusa*, especially for TEs and effector genes (Figure 2B). To understand the origin of this variation, we counted the number of isolates in the accessory nodes of the pangenome graph. The accessory nodes that are present in a low number of isolates (two to six) are most likely due to recent genome expansions that generate unique regions and that are in the process of getting fixed in the population. Conversely, accessory nodes that are present in a high number of isolates (14-18) are most likely caused by deletions of core regions. The total genome size of nodes present in only a low number of isolates is 63% higher than nodes being present in a high number of isolates, suggesting that genome expansions drive the accumulation of unique regions, and are therefore the main origin of the here observed variation between isolates (Figure 2C). In *P. effusa*, genome expansions have been previously associated with the activity of TEs [31], but genes and especially effectors exhibit similar accessory node distribution (Figure 2C), indicating that gene and effector variation is most likely caused by gene copy-number expansions and mutations rather than by deletions.

### Phasing of genomic variation correlates with heterozygosity

Like many other oomycete species, *P. effusa* is a heterozygous, diploid organism [38]. Thus, phasing of the two haplotypes in the genomes is needed to fully account for the observed genomic variation. Due to the obligate biotrophic nature, we do not have access to the parental genomes, and thus it is challenging if not impossible to separate phases (haplotypes) prior to genome assembly. We therefore used a combination of computational approaches based on Nanopore long reads to perform variant calling for each of the 19 genome assemblies and to subsequently phase heterozygous variants. First, variant calling was performed using PEPPER, which phased the Nanopore reads and discovered phased short variants (1-59 bp) [39]. Then, the phased long reads were used with Sniffles2 to discover phased larger insertions and deletions (4-40,000 bp) [40] (Figure 3A). For each isolate, we discovered on average 126,742 heterozygous phased variants, of which the vast majority (98.5%) were identified by PEPPER. Due to their larger size, however, Sniffles2 contributes 72-86% of the variable nucleotides in each genome assembly. Importantly, the large number of phased variants and the length of the Nanopore reads (on average a N50 of 28 kb) allowed for the creation of large, phased blocks (1 Mb on average), up to the full size of complete chromosomes (Figure 3B).

**Figure 3.**
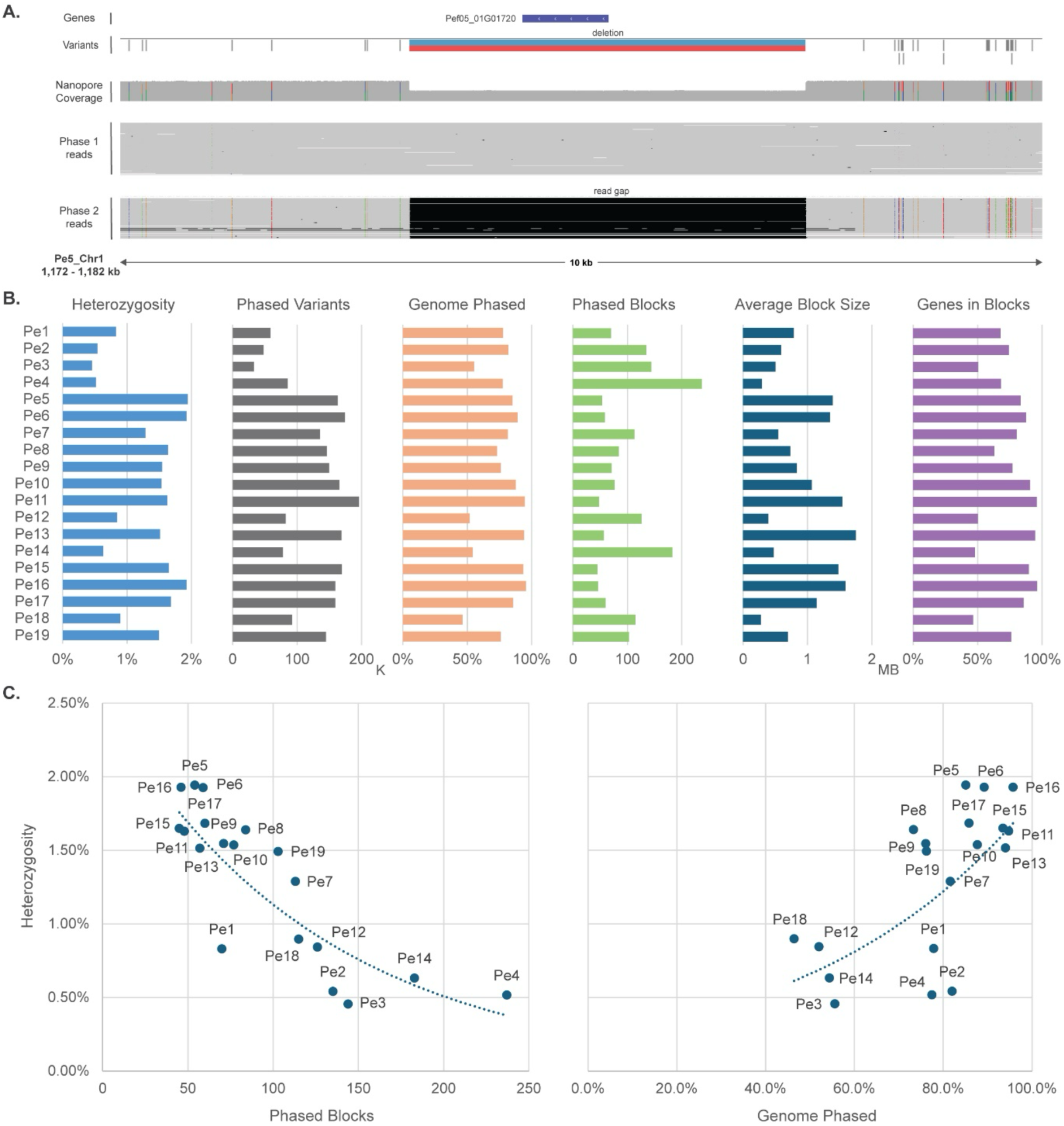
Phasing of *Peronospora effusa* genomes based on variant calling with Nanopore reads. **A.** Example of a heterozygous region in chromosome 1 of *Pe5* visualised with IGV [41]. From top to bottom, the panel shows an annotated gene in the heterozygous region, the variants called from PEPPER with grey lines and the deletion from Sniffles2 with red and blue, the coverage of Nanopore reads, the mapped Nanopore reads that are grouped as phase1, and the Nanopore reads of phase2 where most heterozygous SNPs and the reads gap appear. **B.** Bar plots show the differences in phasing results for the 19 *P. effusa* isolates. It reports the genome-wide heterozygosity, the number of phased variants, the percentage of the genome that is phased, the number of phased blocks, the average size of phased blocks, and the percentage of genes that fully overlap with phased blocks. **C.** Scatterplots show the association between the level of heterozygosity of the 19 isolates and the number of phased blocks (left) or the percentage of the genome that is phased (right).

Heterozygosity, and thus the number of phased variants, varies greatly between the isolates (0.46-1.94% heterozygosity and 32,667-196,278 variants) (Figure 3B). These differences are notable at every aspect of the genome phasing procedure, with highly heterozygous isolates (>1.5%) having less and longer phased blocks (45-77 blocks) that cover most of the genome (>85%), while isolates with lower levels of heterozygosity (<0.9%) have more and smaller sized blocks (115-237 blocks) that cover only about half of the genome (46-55%) (Figure 3B). Consequently, there is a clear negative correlation between the level of heterozygosity and the number of phased blocks and a positive correlation between heterozygosity and the percentage of genome phased (Figure 3C). These observations similarly apply to individual chromosomes, whose heterozygosity varies greatly, both between chromosomes in one isolate as well as between corresponding chromosomes in different isolates. For example, chromosome 13 is almost entirely homozygous in *Pe12* (0.08% heterozygosity), while in *Pe11* it is heterozygous (1.48% heterozygosity). Inversely, chromosome 11 is heterozygous in *Pe12* (2.29% heterozygosity) and mostly homozygous in *Pe11* (0.77% heterozygosity). Thus, the number and coverage of phased blocks vary greatly between chromosomes, and it was not possible to fully phase all chromosomes in even the highly heterozygous genomes. Nevertheless, our approach still enabled us to analyse the heterozygosity of all isolates in depth, and we were able to successfully phase smaller chromosomes that are highly heterozygous.

### *Peronospora effusa* haplotypes reveal multiple evolutionary mechanisms

To compare any genomic region between the 19 isolates, phased or unphased, from a single gene to a full chromosome, we utilised the variation that is directly captured in the pangenome graph. We divided the nodes of the pangenome graph into windows (size specified in each figure) and assign different haplotypes based on the observed variation. In order to not overemphasise small amounts of variation, if all isolates are more than 95% identical in a window, the window was assigned to the core, but if the variation was higher than 5% and the window was only found in one isolate, the window was assigned as a unique genomic region. The windows with accessory variation (>5%) were recursively assigned to isolates based on their genomic relationship; first to isolates from clusters i-iii and then to the isolates that are located closer to the centre of the phylogenetic network (e.g. *Pe11* or *Pe13*) (Figure 1B) (the haplotype order is shown in the corresponding figure legend). We then coloured the nodes in each window based on their assigned haplotype, thus creating haplotype blocks that could be visualised based on their respective position in the pangenome graph (Figure 4A).

**Figure 4.**
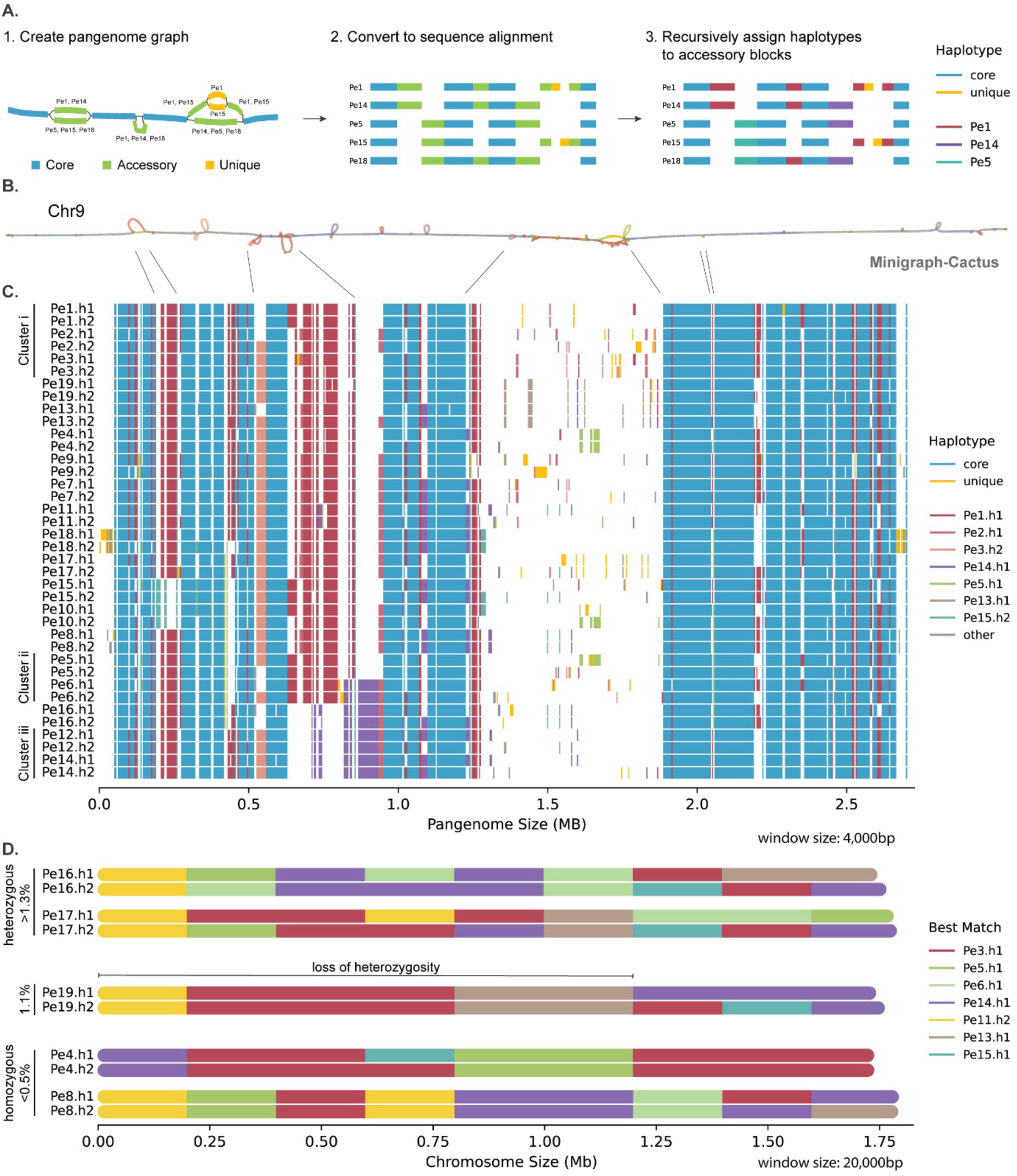
Haplotype comparison of the fully phased chromosome 9 uncovers a variety of evolutionary processes. **A.** Explanation of the here applied approach to assign haplotypes to accessory nodes of the pangenome and subsequent visualization: 1. we create a pangenome graph and parse its variation; 2. the graph is transformed into a sequence alignment for visualisation; 3. the core and unique nodes remain unchanged, while the accessory nodes are recursively assigned to different haplotypes. The order and the colour of the assigned haplotypes is shown in the figure legend. **B.** Pangenome graph of the phased chromosome 9 created with Minigraph-Cactus and visualised with Bandage [36,37]. The nodes of the graph are coloured based on the haplotypes assigned in C. **C.** Alignment of haplotypes of chromosome 9 for all phased *P. effusa* isolates, derived from the pangenome graph in 4 kb windows, as described in A. **D.** For both phases of five isolates, we coloured chromosome 9 based on the best match to a different isolate in a 20 kb window. As targets, we selected the seven isolates that contributed most genomic diversity.

This method was applied to a fully phased chromosome to capture all genomic variation in each isolate, and through their comparison, to discover possible signatures of recombination and differences in the evolution of *P. effusa* isolates. As an example, we focussed on chromosome 9 as it is the smallest *P. effusa* chromosome, is highly conserved, and has a low TE content [31]. Additionally, it has been fully phased for most isolates (85-100% for eleven isolates) and the remaining isolates are either near homozygous (<0.3% heterozygosity for *Pe3*, *Pe4*, *Pe7*, *Pe8*, and *Pe14*) or are overall highly heterozygous, but have lost heterozygosity in large regions of the chromosome (*Pe12*, *Pe18*, and *Pe19*). With the phased chromosome 9 from these 19 isolates, we created a pangenome graph with Minigraph-Cactus [36] (Figure 4B). The pangenome graph of the phased chromosome 9 has 104,800 nodes and 143,918 edges, is highly linear, and 2.7 Mb long: 52% longer than the average size of chromosome 9 and 16.4% longer than the pangenome graph of the unphased chromosome 9 (67,084 nodes and 91,869 edges). Thus, phasing of this chromosome captured additional genomic variation that was not present in the pangenome graph that is solely based on the haploid genome assemblies.

By comparing the different combinations of haplotype blocks on chromosome 9, two main genomic organisations emerged, namely the one observed in cluster i (*Pe1*, *Pe2,* and *Pe3*) and the other one in cluster iii (*Pe12* and *Pe14*). Most haplotype blocks from cluster i that are shared with isolates outside the cluster, are present in *Pe2* and *Pe3*, which suggests that in most cases other isolates have recombined with *Pe2* and *Pe3* rather than the much older *Pe1* isolate. Additional haplotype blocks can be found in a small number of isolates, and these were assigned to *Pe5*, *Pe13*, and *Pe15*. The observed recombination of these haplotype blocks can explain the chromosomal markup of any other isolate, with the addition of few unique blocks, providing strong support of sexual recombination (Figure 4C).

Potential recombination between isolates can be more clearly described by dividing each chromosome into multiple windows and comparing these directly between the isolates, thereby revealing the variation between the phases of different isolates. To this end, we split each chromosome in 20 kb windows and exploited the pangenome graph to assign to each window the best match from a different isolate (Figure 4D). To ease interpretation, we selected one representative isolate from each phylogenetic branch from a hierarchical clustering based on the accessory nodes of the pangenome graph. Note, however, that these are not always identical to the isolates selected when assigning haplotypes (Figure S3). The combination of different matches in a chromosome offers evidence of past recombination events, while the difference between the phases can identify the timeline of those events.

Based on this approach, we selected five distinct examples of isolates to showcase the different processes that contributed to their evolution: *Pe4* is a clear example that demonstrates the occurrence of recombination. It clusters close to isolates from cluster i, but its mitochondrial phylogeny is the same with the isolates in cluster ii. Moreover, *Pe4* also has the accessory chromosome 18, which is not present in isolates from cluster i [29,31]. Evidence of recombination is found in the variation of chromosome 9 of *Pe4* that is most similar to *Pe3*, with some regions matching to *Pe5* and *Pe14* (Figure 4D). Like *Pe4*, *Pe8* is positioned in the middle of the phylogenetic network, has the accessory chromosome 18, and shows clear evidence of recombination as chromosome 9 matches to clusters i, ii, and iii (Figure 1B, 4D). While both *Pe4* and *Pe8* are the result of recombination, they show low levels of heterozygosity throughout the chromosome (0.49% and 0.29%, respectively), suggesting an extended period of asexual reproduction, which is further corroborated by a high number of unique SNPs (*Pe4* and *Pe8* have 7,031 and 9,841 unique SNPs, respectively) (Figure 1B). In contrast, isolates like *Pe16* and *Pe17* have much higher levels of heterozygosity (1.49% and 1.29%, respectively), and display clear evidence of recent recombination events, since their best matches to other isolates differ greatly between their two alleles (Figure 4D). The recent recombination is also evident in the lower number of unique SNPs of *Pe16* (5,740 unique SNPs). However, *Pe17* has 7,730 unique SNPs, suggesting that there might be additional isolates closely related to *Pe17* that are not part of our analyses (Figure 1B). *Pe19* also shows evidence of recombination from distant (*Pe3* and *Pe14*) and closely related isolates (*Pe11*, *Pe13*, and *Pe15*) (Figure 4D), and has 12,528 unique SNPs (Figure 1B). While *Pe19* is heterozygous (1.07%), this variation is in the last third of the chromosome, while the first two thirds show a complete loss of heterozygosity (Figure 4D).

These haploblock recombinations have been showcased here on a single chromosome, but our observations can be generalised for the nearly complete genomes of our collection of isolates. While we could not fully phase the entire genome, the approach of assigning haplotypes based on the best matches between chromosomes can also be applied to the haploid, unphased genomes, which also provided clear evidence of recombination. We observed two groups of isolates: first, isolates that belong to clusters i-iii that share most haplotypes with isolates from within their respective cluster. The second group of isolates display a mosaic pattern of best matches to different isolates (Figure S4). This pattern, with chromosomes being formed by complex combinations of genomic regions from many different lineages, suggests that isolates outside of the three clusters are the result of multiple recent sexual recombination events, which is further corroborated the PHI-test for recombination (p < 0.0001). Our observations from both unphased haploid and phased chromosomes therefore highlight that *P. effusa* isolates evolve by a range of evolutionary processes, which ultimately underlie the rapid emergence of new aggressive *P. effusa* races that can break host resistances.

### Changes in virulence are the result of multiple independent evolutionary adaptations

Variation in effector genes between the 19 *P. effusa* isolates is most likely responsible for resistance breaking in spinach (Figure 1A). Our phased genome assemblies enable us for the first time to explore the full extent of gene and protein variation between *P. effusa* races. We searched for phased variants overlapping with protein-coding genes and integrated them into the gene sequence, thus, when possible, creating two alleles for each gene. For each isolate 2,916 to 35,426 phased variants overlapped with genic regions, affecting between 1,066 up to 5,952 genes, including seven to 101 gene deletions (Table S4). Despite the large number of heterozygous variants that overlapped with genes, for each isolate 94 to 99% of all their proteins are more than 99% identical between the two alleles (Figure S5). Interestingly, while most alleles are highly similar, genes encoding secreted proteins, RXLR and CRN effectors, and especially genes encoding proteins of the same functional group that are physically clustered on the chromosome are enriched for allelic variation (Figure S6).

The presence and absence of genes can be directly inferred from the number of accessory or unique orthogroups, which revealed that on average for each isolate only 12% of all proteins have an ortholog missing in another isolate. Significant genomic variation can also be present in core orthogroups due to mutations, especially when these mutations lead to frame shifts or premature stops. To investigate this, we compared the protein sequence identity in each orthogroup, which revealed that on average 23% of all proteins in each isolate have identical orthologs in both alleles for all *P. effusa* isolates, while half of all proteins have all orthologs in all isolates at least 60% identical in sequence (Figure S7). Importantly, half of all proteins in an isolate miss a similar ortholog (< 60% sequence identity) in at least one allele of at least one isolate, considerably more than the 12% observed just from the gene presence/absence variation. Effector proteins display a similar trend compared with other protein-coding genes, but are more variable (11% identical orthologs, 32% orthologs >60%), while effector proteins that derived from genes that are not part of a physical cluster of effector genes are more conserved (15% identical orthologs, 39% orthologs >60%) (Figure S7). These observations are expected since clustered effector genes are the result of gene copy-number expansions and evolve under positive selection [31].

We have previously shown that unclustered effectors do not typically evolve via gene copy-number expansions, and these genes lack highly similar paralogs in the genome [31]. Consequently, we argued that haplotype variation of single unclustered effectors could be directly connected to the phenotypic differences in virulence between the 19 isolates (Figure 1A). To test this hypothesis, we searched for effectors that can be associated with avirulence of each host resistance gene (Figure 1A), assuming that the effector protein would need to be highly conserved in all avirulent isolates, and inversely, the gene encoding the effector protein should be absent or significantly changed in all the virulent *P. effusa* isolates. For each spinach resistance trait, our approach yielded 20 to 50 effector candidates that were manually inspected, resulting in the identification of four effector candidates we will further discuss to highlight different evolutionary processes that could lead to the emergence of resistance breaking *P. effusa* isolates.

Spinach cultivars with the *RPF3* gene are resistant to ten of the 19 isolates. We identified a single 140 amino acid long effector with a RXLR-EER motif, whose gene is localised in the middle of the large chromosomal arm of chromosome 15, that is highly conserved in twelve isolates (>=98.3% protein sequence identity of both alleles) and absent in seven isolates. This pattern correlates with the *RPF3* resistance, with all seven isolates that lack this effector being virulent on *RPF3* spinach (Figure 5A). Pangenome-enabled comparison of the region around this gene revealed a deletion in a 12.7 kb region that includes three annotated genes and two identical repetitive regions at the start and end of this region, annotated as unknown TEs (Figure 5B). The isolates that lack this genomic region have only one copy of this TE, suggesting that the presence or activity of the TE contributed to the deletion of this region. The analyses of the haplotype blocks of this region reveals that six of the isolates with the deletion are closely related (*Pe7*, *Pe10*, *Pe13*, *Pe15*, *Pe17*, and *Pe18*), possibly sharing a common ancestor. While *Pe2* also contains this deletion, it has a haplotype distinct from these six isolates and is more closely related to isolates without the deletion (*Pe1*, *Pe3*, and *Pe6*) (Figure 5C). These observations collectively indicate that the observed deletions of the region occurred in two separate events.

**Figure 5.**
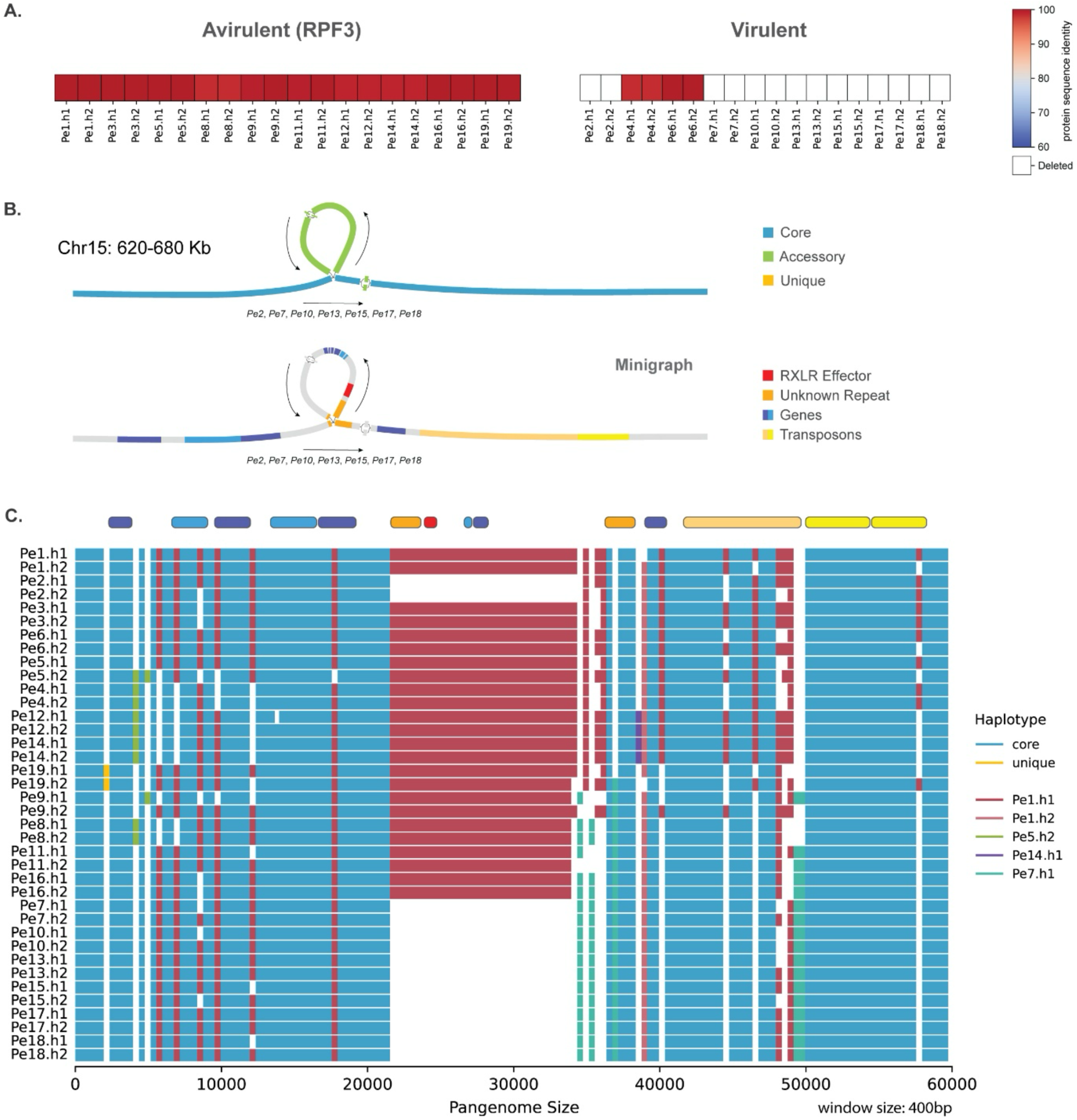
The deletion of a single RXRL effector gene correlates with the resistance breaking of *RPF3*. **Α**. Protein sequence similarity to the longest effector of the Chr15-316 orthogroup, which includes both alleles of all isolates separated by the (a)virulence phenotype on the spinach with the resistance gene *RPF3*. **B.** Pangenome graph of a 60 kb region around the RXRL effector shows the variation in this region and the annotated genes and TEs. **C.** Full alignment of the haplotypes in this region based on the approach explained in Figure 4A, using a 400 bp window. The gene and TE annotations for *Pe1* are visualised on the top.

Spinach cultivars with the *RPF4* gene are resistant to five of the 19 isolates. We identified an effector gene encoding a protein with RXLR-EER and WY motifs, localised on chromosome 15, which is almost identical for 15 alleles (>99.6% protein sequence identity), while the other 23 alleles have only 30.0% identity, 21 of which can be found in isolates that are virulent on *RPF4* spinach (Figure 6A). This is the result of a single nucleotide deletion of the 535^th^ nucleotide of the gene, which causes a frame shift, a downstream premature stop, and thus a truncated protein. As a result, the truncated protein is only 219 amino acids long, much shorter than the 720 amino acids long protein encoded by the allele in avirulent isolates, thus missing the entire C-terminal region of the effector that typically is functional in plant cells [13,18,20]. Notably, this is the only example where a single mutation can be attributed to all variation of this gene between the isolates. This mutation was already present in two heterozygous alleles in otherwise avirulent isolates (*Pe4*.h1 and *Pe15*.h2), indicating that this haplotype was present in the population before the virulent phenotype emerged.

**Figure 6.**
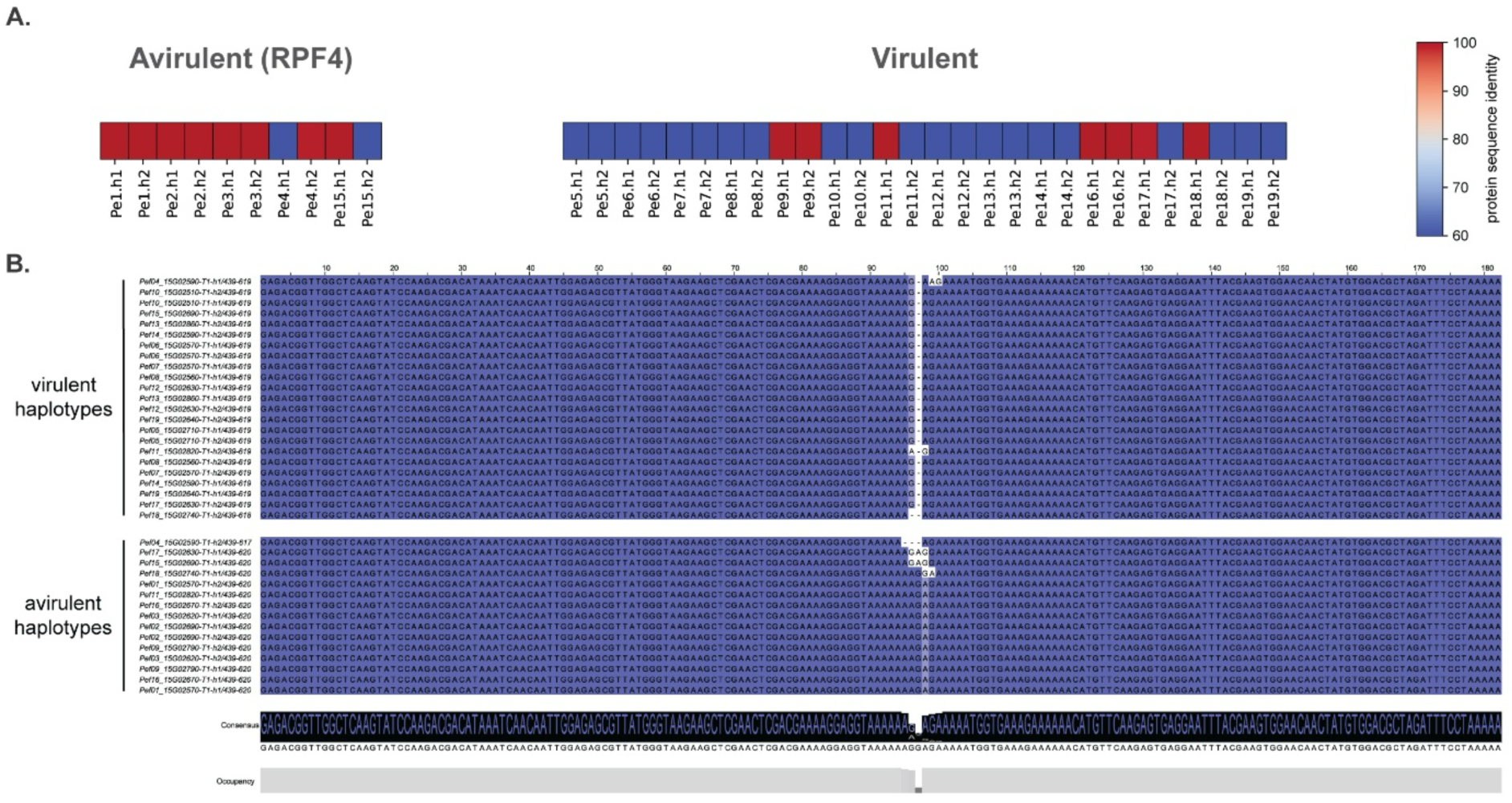
A single mutation in a RXRL effector gene associates with the resistance breaking of *RPF4*. **Α**. Protein sequence similarity to the longest effector of the Chr15-16 orthogroup, which includes both alleles of all isolates, separated by the (a)virulent phenotype against the spinach with the resistance gene *RPF4*. **B.** Nucleotide alignment of a 182 bp long region of the effector genes belonging to Chr15-16 orthogroup separated by isolates with (a)virulence phenotypes.

Virulence on *RPF4* spinach varieties can also be associated with a second effector, 387 amino acid long, with the RLXR-EER and WY motifs, whose encoding genes is localized on chromosome 5. Here, we observed a highly conserved haplotype (>98.2% protein sequence identity) in avirulent isolates and haplotypes in virulent isolates that are either only half the size or completely deleted (Figure 7A). These nonfunctional haplotypes resulted from five separate mutations in the gene, causing early stops in the open reading frame. The appearance of these mutations in the isolates is consistent with the phylogeny of this gene, indicating that there are up to five pseudogenisation events (Figure 7B). Beyond these five mutations, we observed an accumulation of mutations in the gene in the virulent genotype that possibly followed the pseudogenisation events.

**Figure 7.**
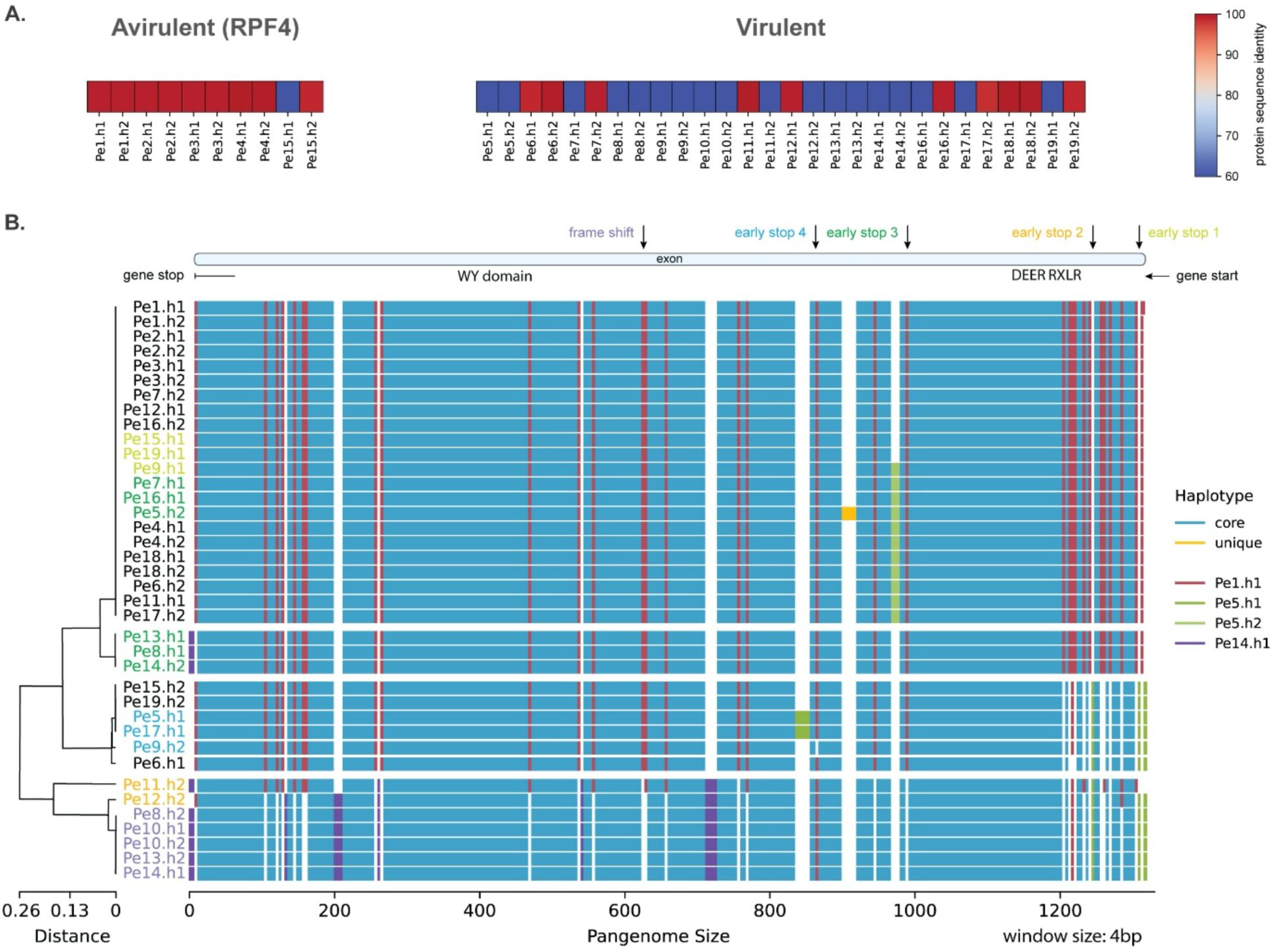
Mutated RXRL effector gene associates with the resistance breaking of *RPF4*. **Α.** Protein sequence similarity to the longest effector of the Chr5-429 orthogroup, which includes both alleles of all isolates, separated by the (a)virulent phenotype against the spinach with the resistance gene *RPF4*. **B.** Full alignment of the haplotypes in this region based on the method on Figure 4A, using a 4 bp window and ordered based on the phylogeny of this region. Isolates are coloured based on the gene mutations or in black for the virulent phenotypes. The gene structure for *Pe1* and the mutated position are visualised on the top.

*RPF11* resistance is broken only by *Pe19* and there is only a single 660 amino acid long effector that is conserved in all other isolates but *Pe19*. The two avirulent haplotypes share at least 97.7% sequence identity, while the virulent haplotypes are 36-56% shorter in length, thus being likely nonfunctional (Figure 8A). These truncations are the result of two different mutations, one SNP creating a premature stop codon that is shared between *Pe19*.h1 and *Pe11*.h1 and one frame shift that causes an early stop codon in *Pe19.h2*. Interestingly, we discovered that the intron is no longer present in *Pe19*.h1 and *Pe11*.h1. These two haplotypes share many haplotype blocks, and thus these likely share a common origin. In contrast, *Pe19*.h2 contains the full intron and has a common origin with the avirulent haplotype 1 (Figure 8B). Consequently, the virulent genotype of *Pe19* has originated most likely from a mutation that already existed in the avirulent isolates and a novel mutation unique to *Pe19*.h2.

**Figure 8.**
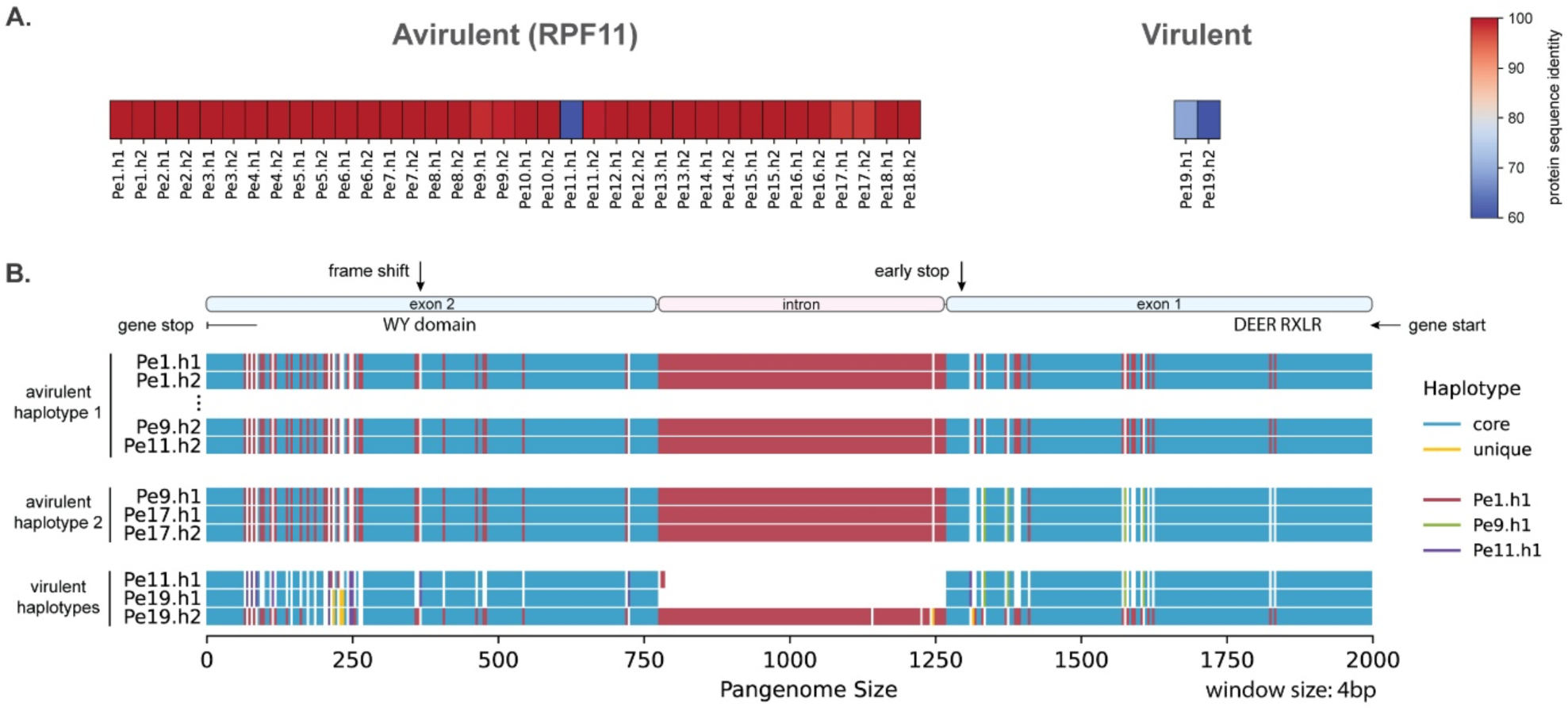
A mutated RXRL effector gene associates with the resistance breaking of *RPF11*. **Α.** Protein sequence similarity to the longest effector of the Chr13-220 orthogroup, which includes both alleles of all isolates, separated by the (a)virulent phenotype against the spinach with the resistance gene *RPF11*. **B.** Full alignment of the haplotypes in this region based on the method on Figure 4A, using a 4 bp window and grouped for virulence against the spinach gene *RPF11*. The gene structure for *Pe1* and the mutated positions are visualised on the top.

## Discussion

Filamentous plant pathogens evolve rapidly to overcome resistances of new crop varieties, often within a few growing seasons [5,6,9]. Resistance of crop varieties is often conferred by qualitative and monogenic effector triggered immunity [7]. Pathogens, in response, adapt their effector repertoire to avoid recognition and maintain virulence [1]. Host NLRs that recognize pathogen effectors have been extensively studied in a few model systems, on fungal pathogens mostly in the genus *Blumeria*, *Magnaporthe*, and *Leptosphaeria*, and in the oomycete pathogens in *Phytophthora* sp. [42]. However, little is known about the diversity of pathogen effector proteins and their evolution, especially in non-model plant pathogens [21,22]. Here we performed, to our knowledge, the largest comparison of oomycete isolates of a single species, by creating a pangenome graph for 19 isolates of the spinach downy mildew *P. effusa* that represent 19 denominated races that are able to break spinach resistances. While the chromosome structure is highly conserved between these isolates, our analysis uncovered an open pangenome, indicating there more variation to be discovered in the *P. effusa* population, due to the extensive variation, mostly caused by copy-number expansions of TEs and effector genes. Variant calling revealed extensive differences in heterozygosity between isolates and between chromosomes within an isolate, which can be explained by the combination of sexual and asexual reproduction and by the loss of heterozygosity in specific genomic regions. Importantly, the fully phased effector repertoires allowed us to pinpoint candidate effectors that are variable between isolates and correlate with the breakage of specific spinach NLR resistance genes. These effector candidates are prime targets for downstream analysis to evaluate their contribution to virulence and function, as well as their potential mode of recognition.

In filamentous fungi, such as in *Verticillium dahliae*, *Fusarium oxysporum*, and *Magnaporthe oryzae*, large-scale chromosomal rearrangements and large accessory regions or chromosomes are often proposed to be important for genetic variation [43–50]. In contrast, the analysis of the 19 *P. effusa* isolate pangenome corroborates that the chromosome structure of *P. effusa* is highly conserved [31]. This conserved chromosome structure is similar to rust fungi, such as previously observed in the genus *Puccinia,* which are also diploids with various degrees of heterozygosity [51–53]. In the absence of large-scale structural variation and accessory chromosomes, smaller mutations (1 bp - 10 kb), allelic variation, and recombination of isolates due to sexual reproduction are the main drivers of evolution [7,51,52].

Oomycetes, like rust fungi, have both sexual and asexual cycles [7,29,54]. Sexual recombination contributes to the evolution of these pathogens by chromosomal admixture between haplotypes, with some races in *Puccinia graminis* formed as a direct result of recombination [51]. Similarly, we provide evidence of recent recombination in five of the 19 *P. effusa* isolates, with seven more showing evidence of past recombination that had been followed by long periods of clonal reproduction. Clonal reproduction can lead to loss of alleles via loss of heterozygosity, which could allow the phenotypic expression of a recessive virulent allele due to the loss of a dominant avirulent allele, leading to isolates that can successfully break resistances [7,51]. We observed loss of heterozygosity in multiple chromosomes of some *P. effusa* isolates, for example in chromosome 9 in *Pe19* (Figure 4C), which may result from genetic bottlenecks or selective sweeps. These cases indicate that *P. effusa* often evolves clonally under strong evolutionary pressure, probably in the conditions created on commercial fields with monocultures of resistant spinach varieties.

Oomycete RXLR effectors, most commonly in *Phytophthora* species, have been shown to suppress plant immunity, like AVR1 and PITG20303 of *P. infestans*, PpE18 of *P. parasitica*, PcAvr3a12 of *P. capsici*, and RxLR50253 of *Plasmopara viticola* [55–59]. RXLR effectors also trigger plant immunity, like AVR1 of *P. infestans* in R1 potato plants, multiple effectors with a WY domain in *Bremia lactucae* in lettuce, and Avr1b and AvrNb of *Phytophthora sojae* in *Nicotiana benthamiana* [20,60–64]. Here we investigated effector variation in *P. effusa* isolates and its correlation with their virulence in different spinach varieties. In most of these cases, the candidate effector genes in the avirulent isolate are highly conserved, while the alleles in virulent isolates accumulate non-synonymous mutations that often result in frame shifts. The most prevalent mechanism that has been shown to avoid recognition are effector gene deletions or point mutations, as shown in *P. infestans* where the virulent allele of *PiAvr4* encodes a truncated protein caused by frame shift, or in *Plasmopara viticola* were multiple RXLR genes have been deleted as a result of structural variations [65,66]. Similarly, we here identify candidate effectors that can be associated with resistance breaking due to gene deletions and truncation of proteins. Nevertheless, we could not discover a case where a single event can be attributed to the virulent phenotype of all isolates, but resistance breaking could be linked to multiple independent evolutionary events. Additionally, we also observed isolates that have both alleles that we have linked to the avirulent effector genotype, thus the absence of the effector protein needs to be achieved via an alternative mechanism, possibly through repression of expression via gene silencing (e.g. *Pe4* and *Pe6* for *RPF3*) (Figure 5A). Epigenetic gene silencing to repress effector expression and avoid recognition in resistant crops has been previously demonstrated in *Avr1b* and *Avr3a* of *P. sojae* [67,68]. Our observations suggest that both epigenetic and ‘conventional’ mutational processes jointly contribute to the resistance breaking of spinach. However, the complex evolution and the extensive genomic variation between the isolates make it challenging to pinpoint the exact mutations that caused resistance breaking. To improve our ability to detect avirulence effectors and understand their evolution, the study of a much larger number of isolates with known (a)virulence phenotypes would be necessary. Alternatively, a small number of additional isolates could already provide better resolution for specific resistances by selecting multiple isolates with identical phenotypes or by re-sequencing isolates for which a new phenotype has emerged after their initial isolation. These results can then be used as the starting point for downstream functional analysis at the molecular level to test the avirulence of effectors in plant tissue [69].

## Conclusions

Like many other downy mildews, the rapid evolution of *P. effusa* leads to the emergence of multiple resistance breaking isolates, often in a single cultural season [24].Together with its obligate biotrophic nature, this poses a significant challenge to isolate, phenotype, maintain, and study this devastating pathogen [12,38,70]. The recent advancements in sequencing and in comparative genomics such as the here applied pangenome graphs now start to enable us to uncover the genomes of these pathogens, to differentiate between isolates and to link this variation to phenotypic differences, thereby providing effectors as prime candidates for future experimentations. The computational approach described here, will continue to uncover detailed mechanisms of the rapid evolution of these pathogens, perhaps leading to predictions of the upcoming emergence of new phenotypes.

## Methods

### *Peronospora effusa* infection on soil-grown spinach and spore isolation

Spinach plants were grown in potting soil (Primasta, Netherlands) under long-day conditions (16-hour light, 21 °C). Two to three weeks post-germination, plants were inoculated with *P. effusa* by spraying them with a spore suspension in water using a spray gun. After inoculation, plants were kept under 9-hour light at 16 °C in humidified trays, where lids were sprayed with water and kept covered. Vents were opened after 24 hours, and lids were re-sprayed and resealed 7–10 days post-inoculation to maintain humidity and promote *P. effusa* sporulation.

For spore collection, sporulating leaves were placed in a glass bottle with tap water and shaken to release spores. The suspension was filtered through a 50-μm nylon mesh (Merck Millipore, USA) to remove large debris and then through an 11-μm nylon mesh using a vacuum pump to remove smaller contaminants. Spores retained on the filter were washed, scraped off, and stored at −80 °C for Oxford Nanopore sequencing.

### High-molecular weight DNA extraction protocol

High-molecular-weight (HMW) DNA was isolated from *P. effusa* spores by grinding them into a fine powder in liquid nitrogen with 0.17–0.18 mm glass beads. The powdered spores were washed with cold sorbitol solution (100 mM Tris-HCl, 5 mM EDTA, 0.35 M sorbitol, 1% PVP-40, 1% β-mercaptoethanol, pH 8.0). Lysis was performed in extraction buffer (1.25 M NaCl, 200 mM Tris-HCl pH 8.5, 25 mM EDTA pH 8.0, 3% CTAB, 2% PVP-40, 1% β-mercaptoethanol) containing proteinase K and RNase A, incubated at 65 °C for 60 minutes with gentle inversion. Debris was pelleted by centrifugation. HMW DNA was purified using phenol/chloroform/isoamyl alcohol (IAA) and chloroform/IAA extractions, followed by an additional RNase treatment, purification with phenol/chloroform/IAA and chloroform/IAA, and isopropanol precipitation. DNA concentration and integrity were assessed using Nanodrop, Qubit, and Tapestation.

### Genome sequencing using Oxford Nanopore

We obtained long-read sequencing data for 13 *P. effusa* isolates with Oxford Nanopore sequencing technology (Oxford Nanopore, UK) at the USEQ sequencing facility (the Netherlands). We used a Nanopore PromethION flowcell (R9.4.1) for real-time sequencing and base-calling of the raw sequencing data was performed using Guppy (version 4.4.2; default settings).

### Genome assembly

To produce chromosome-level genome assemblies of thirteen *P. effusa* isolates, we used the long-read Oxford Nanopore sequencing data. The reads were corrected, trimmed, and assembled using Canu (version 2.3) [71] with the following command:

~~~
canu -nanopore ${input_reads} genomeSize=58M -d ${output_dir} -p ${isolate_name}
corOutCoverage=40 mhapMemory=100g corMhapFilterThreshold=0.0000000002 mhapBlockSize=500
ovlMerThreshold=500 corMhapOptions=”--threshold 0.80 --num-hashes 512 --num-min-matches 3
--ordered-sketch-size 1000 --ordered-kmer-size 14 --min-olap-length 800 --repeat-idf-scale 50”
~~~

### Scaffolding assemblies and closing gaps

The assemblies were scaffolded to full chromosomes with ragtag scaffold (v. 2.1.0, default settings) [72] using as reference the most closely related genome assembly from the previous assembled *P. effusa* isolates (*Pe1*, *Pe5*, *Pe11*, *Pe14*, and *Pe16*) [31]. Gaps in the scaffolded chromosomes were closed with FinisherSC (version 2.1; default settings) [73], and the scaffolded assemblies were corrected for single nucleotide polymorphisms using Illumina short-reads with four rounds of Pilon (version 1.23; --diploid, --fixbases) [74].

### Transposable element and genome annotation

The combined *P. effusa* TE library created from *Pe1*, *Pe5*, *Pe11*, *Pe14*, and *Pe16* [31] was used to annotate and soft-mask the genomes using RepeatMasker (version 4.1.2 -e rmblast -xsmall -s -nolow) [75]. The soft-masked genomes and RNAseq short-read data were used for structural gene prediction and functional annotation with the funannotate pipeline (version 1.8.7) [76] as described previously [31].

### Secretome and effector prediction

To detect secreted proteins and effector candidates, we applied a previously published approach [31]. In short, we used the genes annotated by funannotate and additional open reading frames, encoding at least 70 amino acids, to predict the secretome using the Predector pipeline (v. 1.2.6, default settings) [77]. The secreted proteins were then screened to detect the presence of the conserved motifs described in RXLR and Crinkler oomycete effectors in the canonical and divergent forms. The search was performed using regular expression with the EffectR package for R [78] and sequence profile searches using HMMER v3.3 [79].

### Pangenome graphs and common annotation

A pangenome graph was built per chromosome and then merged in a final pangenome graph following the Minigraph-Cactus Pangenome Pipeline, HPRC Graph (step-by-step): Splitting by Chromosome (version 2.6.4, --filter 0 --vcf full --gfa full) [36]. This results in one, unfiltered pangenome graph of all 17 core chromosomes of all 19 isolates that was used in the downstream analysis. Pangenome graphs from Minigraph were visualized with bandage (version 0.8.1) [37].

The hal output of the pangenome graph, the gene annotation, the annotated protein sequences, and the RNAseq coverage for each *P. effusa* isolate were used as input to collectively reannotate genes with the Comparative-Annotation-Toolkit (version 2.2.1, --augustus --augustus-cgp --assembly-hub -- filter-overlapping-genes) [80]. To filter ORFs of unknown origin, we removed from the annotation genes that were not characterized as protein-coding (gene_biotype=protein_coding) or had no “alternative_source_transcripts” in the gff line of the tRNA.

### Synteny-based gene orthogroups

Synteny based gene orthogroups were created based on our previously published approach [31]. In short, the gfa file of pangenome graph was parsed and annotated with the genes of each isolate. The genes were then linked in orthogroup based on their syntenic localization along the graph. In total, this resulted in 15,739 orthogroups with a single gene for each isolate represented in each orthogroup.

### Saturation plots

Saturation plots are created based on our previously published approach [31]. Briefly, the variation between the isolates was visualised by making all the possible comparison for combinations of one to all 19 isolates. We characterised each count based on the number of isolates represented, as core (all isolates in the comparison), unique (only one isolate for comparisons of two isolates or more), or accessory. The line is drawn on the mean for each combination (one to 19) and the range of all calculations are shown by the shadow behind each line.

### Haplotype assignment

Each variable node, i.e., each node that is not present in at least one isolate, in the parsed pangenome file was assigned a haplotype. Nodes were assigned as a unique if they only occurred in one isolate. The rest of the nodes were recursively assigned to a haplotype, based on the isolates represented in the node. For a given window size, we calculated the haplotype that is most abundant by length. If the total length of that haplotype is above 5% of the windows size, the whole window is labelled to belong to that haplotype. Otherwise, the window is characterised as core. The alignment of the region is visualised in python using pandas and matplotlib [81].

### Variant calling with Nanopore reads

Nanopore read of each isolate were mapped to the respective genome assemblies using minimap2 (v. 2.21, -ax map-ont) [82]. Variant calling of short variants and the phasing of the nanopore reads was performed with the PEPPER-Margin-DeepVariant pipeline (v. 0.8) [39] with the following command:

~~~
apptainer exec --bind ${home_dir} pepper_deepvariant_r0.8.sif run_pepper_margin_deepvariant
call_variant -b “${nanopore_bam}” -f “${ref_fasta}” -p “${isolate}_PEPPER_Margin_DeepVariant”
-t “${threads}” -o “${output_dir}” --ont_r9_guppy5_sup
--phased_output --pepper_min_mapq 10 --dv_min_mapping_quality 10
--margin_haplotag_model allParams.haplotag.ont-r94g422.json
--margin_phase_model allParams.phase_vcf.ont.json
~~~

The bam file of phased nanopore reads was then used to perform variant calling for phased long variants using Sniffles2 (v. 2.3.3, --phase). The phased insertions and deletions discovered by Sniffles2 were merged with the phased short variants from PEPPER by giving priority to a deletion when it overlapped with other SNPs in the same phase.

### Phasing of genomes and genes

The merged vcf file of phased variation was used to create two copies of each genome assembly, one for each phase with bcftools consensus (v. 1.16, -e ’ALT∼”<.*>”’) [83]. Individual chromosome of the phased copies from the genome assemblies were used to create phased pangenome graphs as described above, by adding the haplotype after the isolate name (e.g. *Pe5*.1 and *Pe5*.2) [36].

### Effector comparison

For each effector orthogroup, we calculated the protein sequence similarity of all proteins compared to the longest allele. To narrow down our search for effector linked to avirulence, we filtered for orthogroups that combine the following characteristics: i) all avirulent isolates have at least one allele with 85% protein sequence similarity; ii) the group of virulent isolates have at least one isolate with both alleles with less than 85% sequence similarity. Heatmaps of the outcome were visualised in python using matplotlib and seaborn [81,84].

### Splits tree

We performed variant calling, using the genome assembly of *Pe1* as a reference, and the short-read data of 19 *P. effusa* isolates. The short reads were aligned using bwa-mem2 (version 2.2.1, default settings) [85] and a joint VCF file was generated with both variant and invariant sites with GATK (version 4.4.0.0, GenotypeGVCFs -all-sites) [86]. The single nucleotide variants were transformed into a distance matrix with PGDSpider (version 2.1.1.5) [87], which was then used to construct a decomposition network using the Neighbor-Net algorithm with SplitsTree (version 4.17.0) [88]. We calculated the branch confidence of the network using 1,000 bootstrap replicates.

## Declarations

### Data availability

For this study, we sequenced isolates of 19 denominated races of *P. effusa*. Cultures of these can be requested for research from Naktuinbouw, the Netherlands [89] (Table S5). The raw sequence data and genome assemblies and annotations generated in this study are available on NCBI under BioProject PRJNA772192. The genome assemblies, gene and repeat annotation, repeat library, effector clustering, gene variation, and pangenome graph are available on Zenodo (DOI 10.5281/zenodo.15490946). The code used in this publication is available on GitHub: https://github.com/TeamMGE/Skiadas2025_pangenome_haplotypes

### Author Contributions

PS and MFS conceived and designed the data analysis, JE and MNM generated the Nanopore sequencing data. PS performed the data analysis, visualised the results, and drafted the manuscript. MFS, RDJ, and GVDA acquired funding for this project, supervised the research, and reviewed the manuscript. All authors approved the final submission.

## Acknowledgments

We acknowledge the Utrecht Sequencing Facility (USEQ) for providing sequencing service and data. USEQ is subsidized by the University Medical Center Utrecht and The Netherlands X-omics Initiative (NWO project 184.034.019). We thank Jan Kees van Amerongen for the invaluable support and the maintenance of the computational infrastructure.

## Supplementary Figures

**Figure S1.**
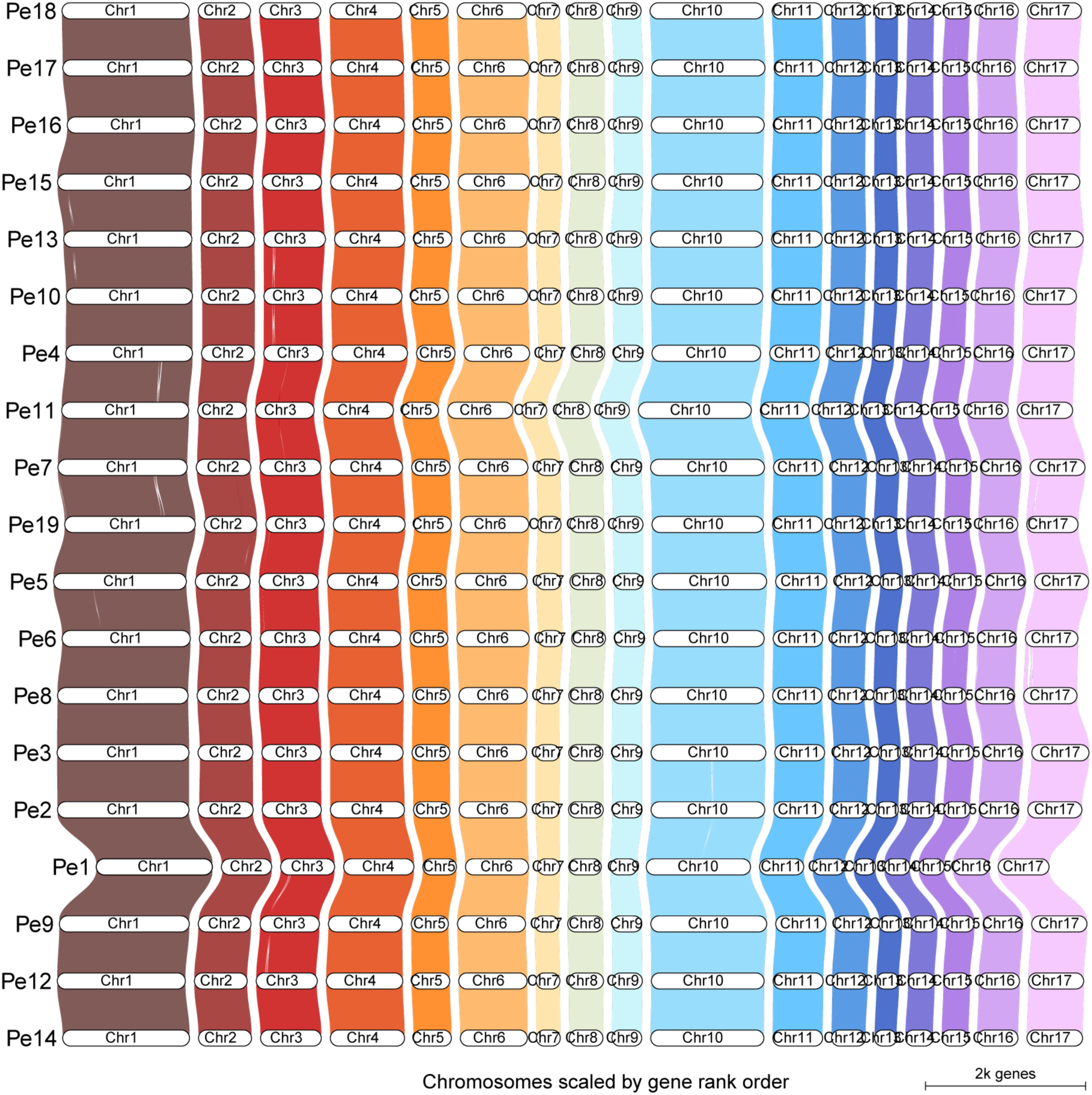
Whole-genome alignments of genomes based on sequence similarity and on relative position of protein-coding genes. Comparison of our 19 chromosome-level genome assemblies for *P. effusa* revealing highly conserved chromosome structure.

**Figure S2.**
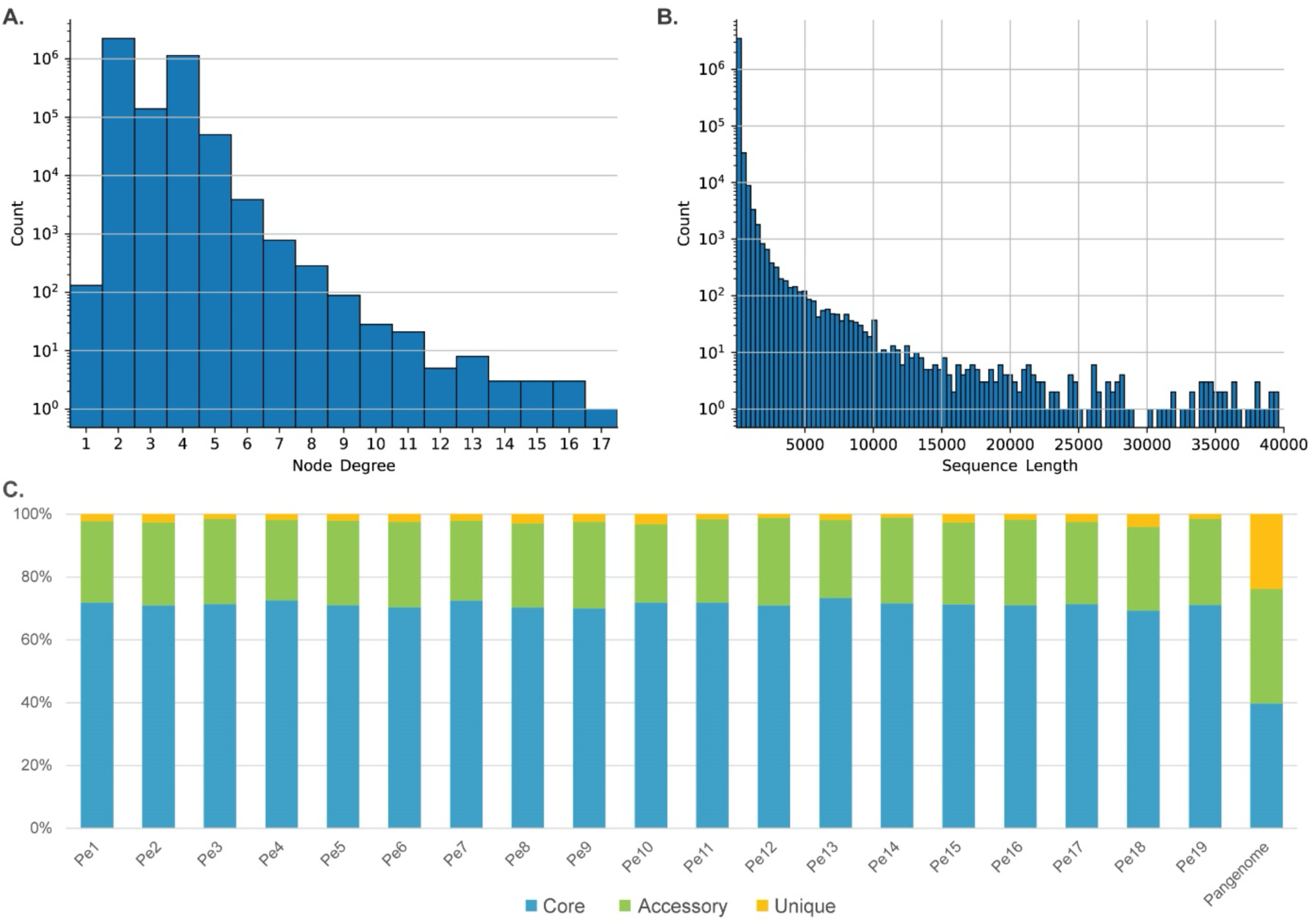
The structure and variation of the pangenome graph of 19 *Peronospora effusa* isolates. **A.** Histogram of the node degree, i.e. the number of connections, of each node of the pangenome graph. **B.** Histogram of the sequence length of each node of the pangenome graph up to a max length of 40 kb. **C.** Bar plot of the percentage of the genome size that is core, accessory, or unique for each isolate and the pangenome.

**Figure S3.**
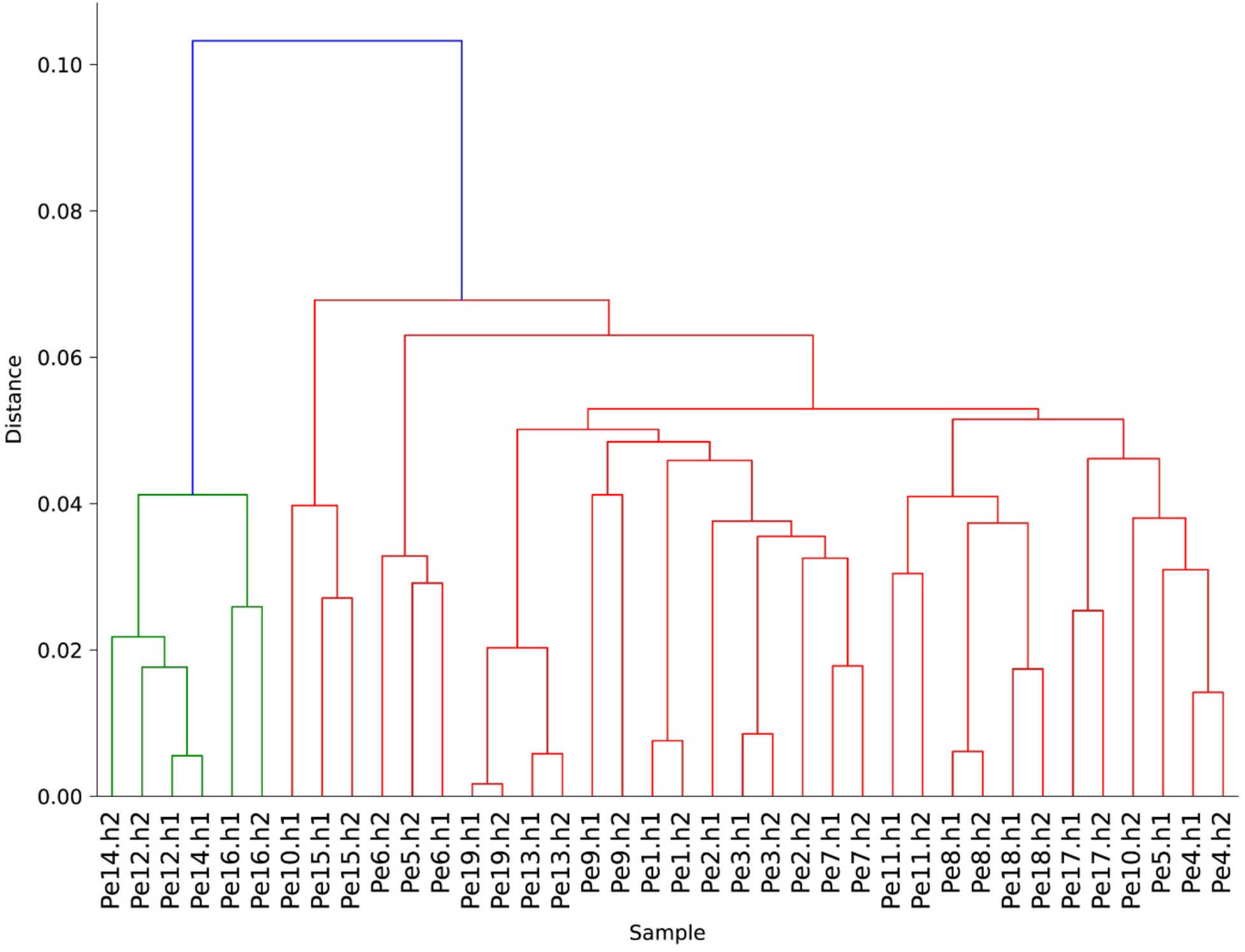
Phylogeny of *Peronospora effusa* haplotypes bases on chromosome 9. Dendrogram based on the hierarchical clustering of the accessory nodes of the phased chromosome 9 from 19 *P. effusa* isolates.

**Figure S4.**
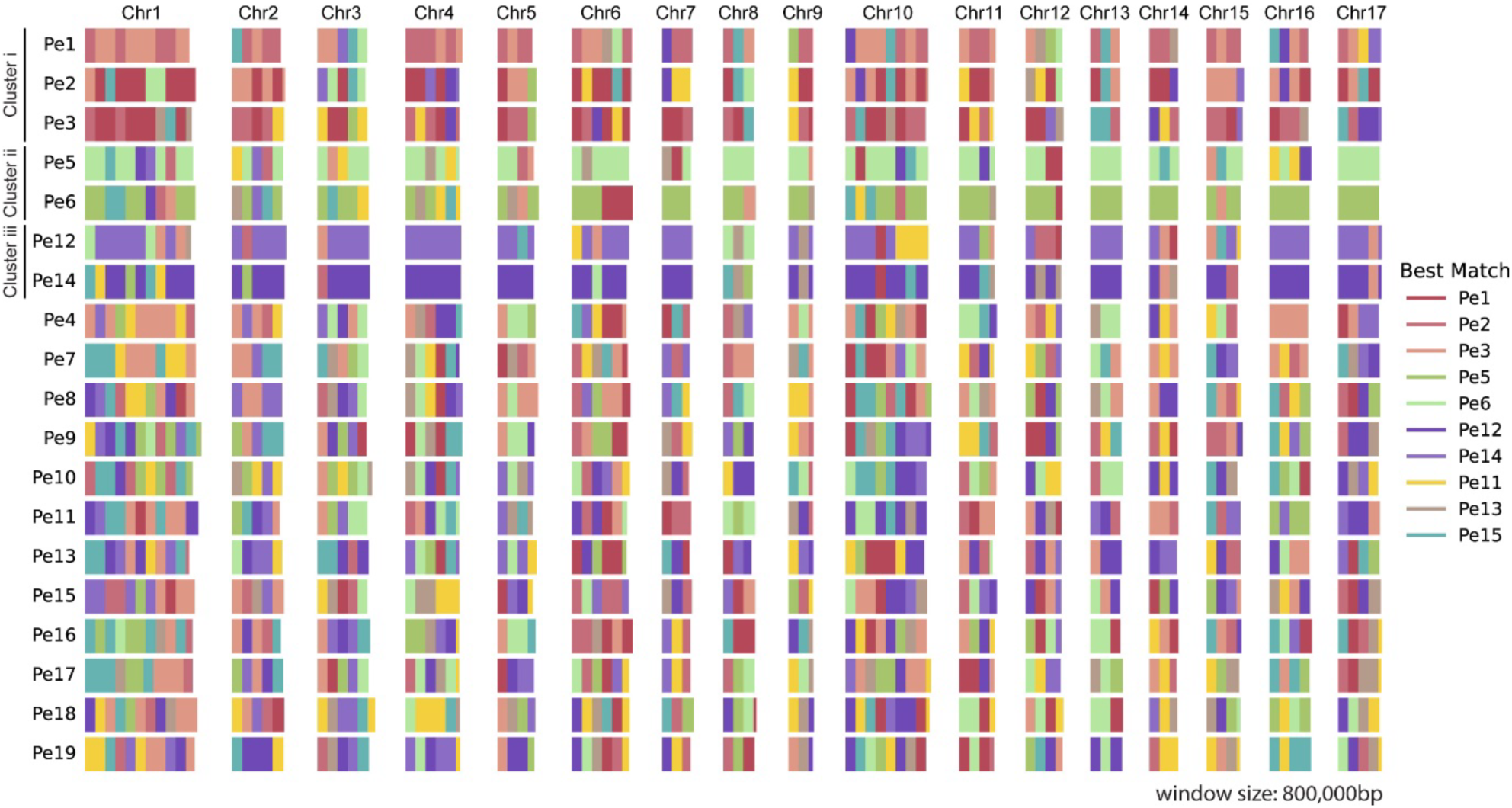
The similarity of the 17 chromosomes of 19 *Peronospora effusa* isolates. The haploid chromosomes are coloured based on the best match to a different isolate. The chromosomes where split in an 800 kb window to offer an overview of all the chromosomes in a single figure. As targets we selected the ten isolates with most contribution to the genomic diversity based on the method described in Figure 4A.

**Figure S5.**
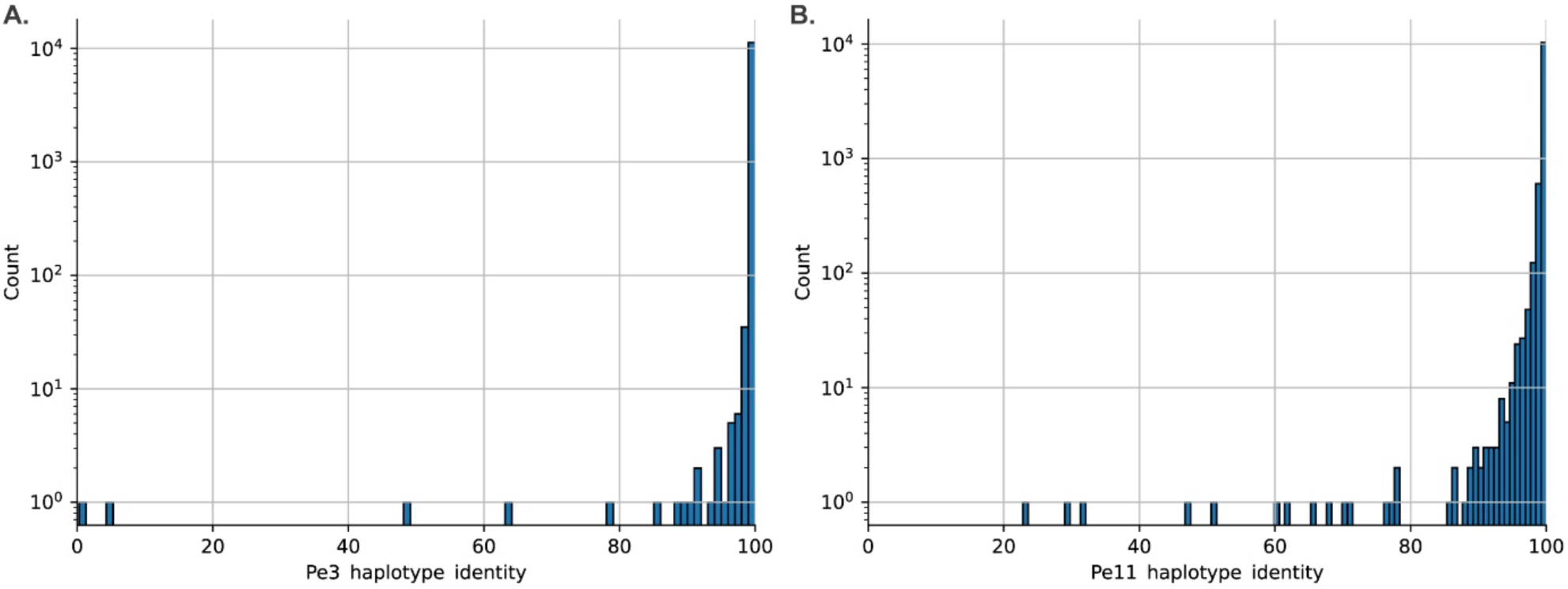
Histograms of the protein sequence identity of haplotypes after phasing. **A.** Histogram of *Pe3* with the least number of phased variants applied to genes. **B.** Histogram of *Pe11* with the most phased variants applied to genes.

**Figure S6.**
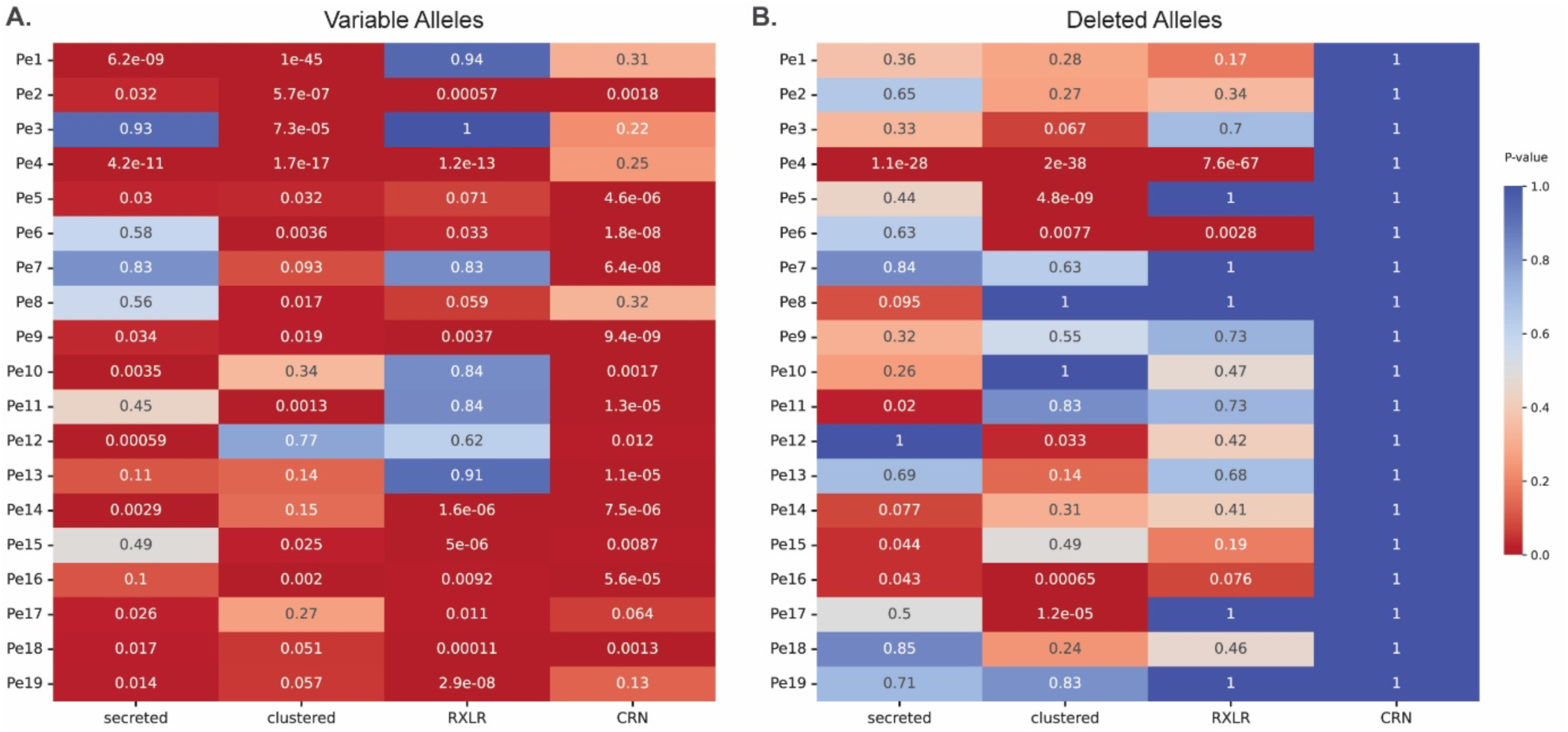
Enrichment of subgroups of protein for variation between alleles. **A.** Heatmap of enrichment for any sequence variation between protein alleles. **B.** Heatmap of enrichment for protein deletion between protein alleles.

**Figure S7.**
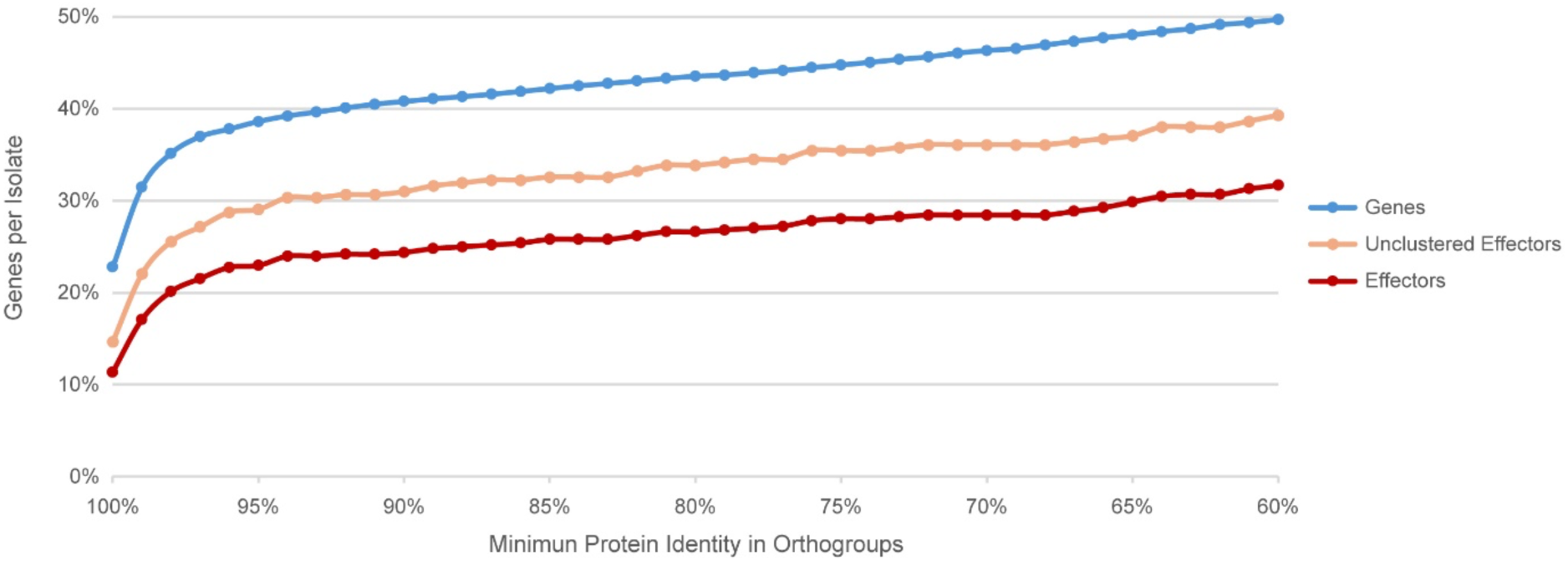
Comparison of the protein sequence similarity between 19 *P. effusa* isolates for different groups of genes. Line plots for the cumulative number of genes that have orthologs in all 19 isolates with a minimum protein identity. Plots are made for all genes, effectors, and effector outside physical clusters.

